# Individuals matter: habitat factors and plant traits shape individual-level pollinator interactions in a semi-arid landscape

**DOI:** 10.1101/2025.07.26.666931

**Authors:** Diana Michael, Kunjan Joshi, Shivani Krishna

**Affiliations:** Department of Biology, Trivedi School of Biosciences, Ashoka University, Sonepat, India

**Keywords:** Flower production, Intraspecific variation, Microhabitat heterogeneity, Resource trade-off, Soil moisture

## Abstract

**Background:** Plant-pollinator interactions are vital for understanding ecological processes influencing reproductive success in plant communities. While species-level pollinator interactions are important for predicting community stability, it remains equally crucial to understand individual-level interactions of keystone species in the community. This study examined the role of habitat factors and floral traits in shaping pollinator interactions at the individual plant scale of *Maytenus senegalensis*, a dominant native species in the semi-arid Aravalli Hills. We measured flower production, nectar sugar concentration, flower diameter, and external factors such as percentage soil moisture, distance to habitat edge, and density of co-flowering conspecifics to assess their impact on pollinator interactions and reproductive success.

**Results:** We found significant variation in reproductive investment in the form of flower production and a trade-off with reward quality, where plants with higher flower production were found to have a lower nectar sugar concentration. Higher flower production negatively influenced reproductive success, suggesting the likelihood of increased within-plant visitation. *Eristalinus* and *Apis* were the dominant pollinator genera, and overall, Dipterans were found to play a critical role in maintaining the network stability. The presence of flowering conspecific plants in the neighborhood reduced the pollen deposition, suggesting competitive interactions. Moreover, individual plants were found to show some amount of specialization in their interaction niches. We predict that this could lead to further divergence of interaction niches due to pollinator-mediated competition. Any perturbation to interactions of plants with a high degree of pollinator connectance was found to disproportionately influence the network.

**Conclusions:** Overall, our results link microhabitat (soil moisture) and neighborhood context to individual interaction niches, demonstrating that allocation trade-offs and conspecific competition jointly shape pollination and fitness. In semi-arid systems, which are undergoing considerable anthropogenic and climatic changes, our study provides insights into individual pollinator interaction niches and the role of microhabitat factors in species persistence within a community.

## Introduction

Plant–pollinator interactions are fundamental ecological relationships that facilitate pollen transfer and reproductive success in flowering plants. These interactions are often driven by the availability and quality of floral resources, which influence pollinator visitation patterns and, consequently, fruit and seed set [1]. The stability of these interactions is crucial for maintaining plant communities [2, 3]. In arid and semi-arid communities, anthropogenic pressures coupled with climatic changes are disrupting these relationships [4, 5]. One of the ways these interactions are being affected is via reduced reproductive investment. Montane plant communities exposed to warmer and drier conditions were found to show clear signs of drought stress, including early senescence and diminished allocation to both reproductive structures and vegetative growth [6]. Studies have demonstrated that climate warming negatively affects plant flowering times, disrupting plant-pollinator interactions and, thereby, reproductive success [7, 8].

Variations in soil moisture influenced reproductive investment directly while also modifying the microhabitat for the organisms that interact with plants [9]. Experimental studies revealed that drought-stressed plants produced significantly fewer flowers and reduced nectar per flower, while pollen quantity per flower remained unchanged. Additionally, nectar from drought-stressed plants exhibited a lower sucrose-to-total sugar ratio, although the overall sugar concentration was unaffected [10]. Plants, being sessile and highly responsive to microhabitat conditions, contrast with the mobility and predominantly generalist feeding habits of pollinators. As a result, plants are expected to exhibit greater spatial variability in their traits than pollinators [11]. Despite apparent uniformity in the landscape, both flowers and pollinators demonstrate significant heterogeneity, creating a mosaic of distinct local communities. Therefore, examining these variations at the level of individuals is critical to accurately understand and predict interaction patterns, and reproductive outcomes. Ignoring individual heterogeneity may bias estimates of interaction structures, and obscure fine-scale reproductive hubs that are natural targets for management and conservation actions [12, 13].

Plant-pollinator interaction studies are typically focussed on how interaction patterns vary between species, while intraspecies interactions remain largely unexplored [14]. Few studies have concentrated on plant-pollinator interactions at the individual plant level [15, 16]. Additionally, studies integrating abiotic and biotic conditions to explain these individual-level plant-pollinator interactions are limited [13, 17]. Individual plants within populations exhibit phenotypic variation, such as differences in plant and flower size or flowering phenology, which can influence their pollination niche by attracting different pollinator guilds. Plants with similar phenotypes tend to interact with comparable pollinator assemblages [18]. Additionally, the spatial location of a plant within the population could influence its pollinator niche, as it is influenced by microhabitat conditions, neighborhood effects (local plant community composition, competition), which shape the context-dependent outcomes of pollinator interactions [19]. As individual plants respond differently to these variations, investigating these patterns at the individual level allows us to understand how variation among plants contributes to overall pollination success and reproductive output at the community level [20, 21].

The preferences of pollinators vary with floral traits, resulting in diverse interaction niches. For example, bee visitation was influenced by pollen production and flower height, and fly visitation was associated with flower size and number [22, 23]. Notably, bees favoured plants with higher pollen production, while flies were drawn to larger flowers with more nectar [24]. Furthermore, the neighborhood of the plant also plays an important role in pollinator behaviour and interactions. Plants near more attractive species often benefit from ‘spillover’ effects, where pollinators attracted to the more prominent species also visit the less attractive species [25]. A similar pattern can be expected at the population level, which could increase the visitation rates for the less attractive plants, even if their floral traits alone would not typically draw as much attention. However, when plant individuals compete for pollinator resources, the dynamics can shift. Increased plant density and abundance can also lead to competitive interactions, diluting the effectiveness of visitation [26]. Such pollinator-mediated, plant-plant interactions can range from facilitative to competitive in nature, with pollinator behaviors (e.g., visitation rates and foraging patterns) influencing pollen deposition and reproductive success. Recent studies have shown that individual plant traits, such as floral position and nectar concentration, can impact pollinator foraging patterns [27, 28], emphasizing the role of floral traits in determining reproductive success.

Here, we examined individual-level variations among plants in pollinator interactions, their stability, and consequences for reproductive success in the semi-arid shrub *Maytenus senegalensis* (Celastraceae; Celastrales). To connect microhabitat level variations to interaction outcomes, we quantified soil moisture around each focal plant, floral investment (flower production), floral traits (e.g., flower diameter), and floral rewards (nectar sugar concentration), along with the density of neighboring flowering conspecifics. For each individual plant, we recorded pollinator visitation, measured conspecific pollen loads, and estimated fruit set. We assessed these links using the framework of network theory to examine the degree of specialization, niche overlap among plants, and robustness to perturbations, and whether these links relate to the measured traits (Fig. 1).

**Fig. 1.**
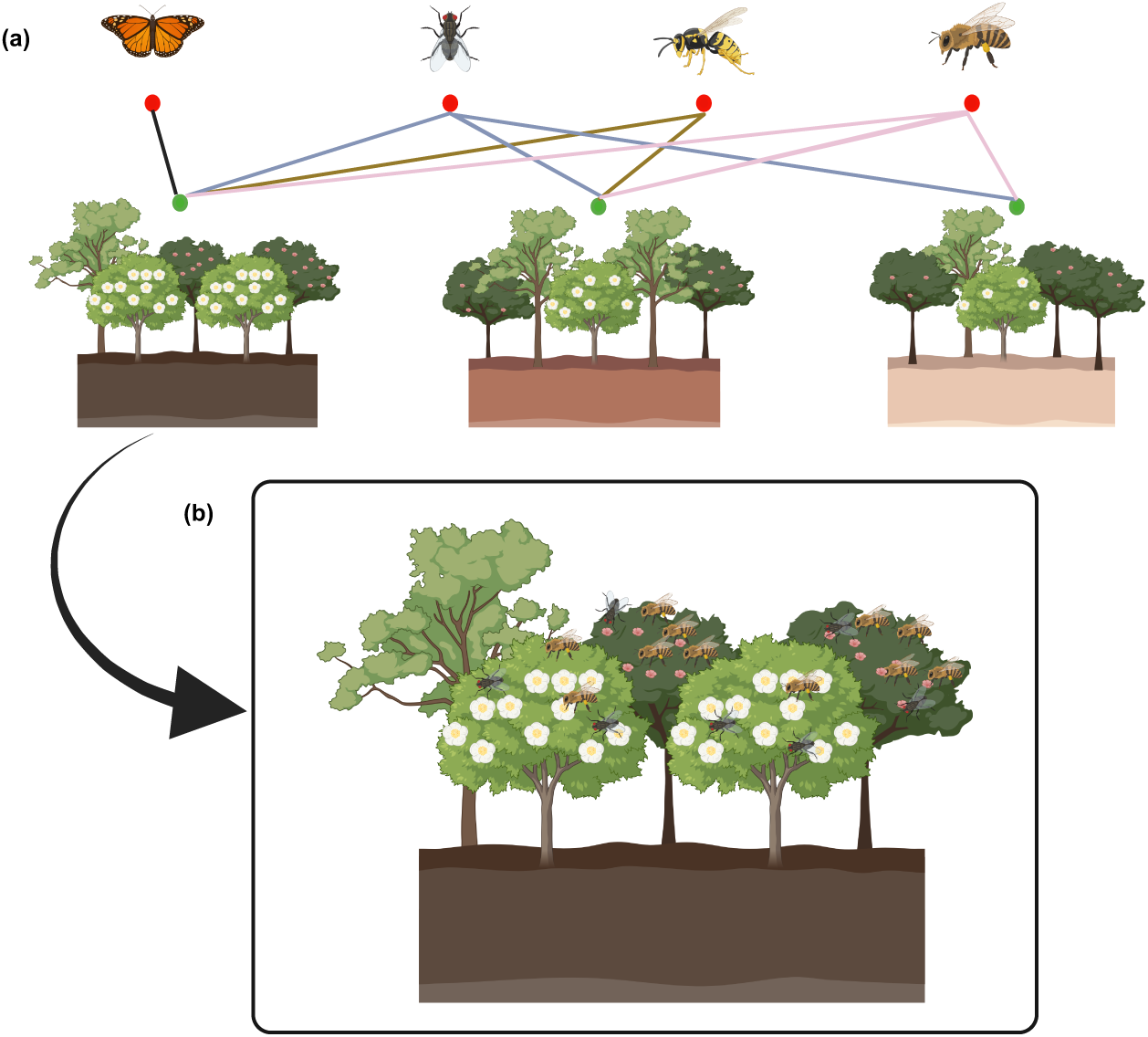
Conceptual overview linking microhabitat heterogeneity to plant–pollinator interactions and fitness in *Maytenus senegalensis.* (a) Microhabitat variation illustrated as a soil-moisture gradient (darker soil = wetter), is hypothesized to alter floral traits and reproductive investment (e.g., flower production) as well as local floral diversity, thereby shaping pollinator-mediated interactions. These interactions are depicted as a bipartite network with pollinators (red nodes) linked to individual plants (green); thicker links indicate stronger visitation. (b) Within a microhabitat, focal plants co-occur with conspecific and heterospecific neighbors and may experience facilitation or competition for pollinator access, depending on neighborhood composition and density. Arrows summarize expected pathways from microhabitat to traits to visitation, culminating in reproductive success (fruit set and conspecific pollen receipt), which we evaluate with individual-level data. Accordingly, we predicted that soil moisture and conspecific flowering density would modulate visitation and fruit set.

Using the backdrop of resource allocation theory [29] and competition-facilitation dynamics [30] in water-limited systems, we made the following predictions: (i) Higher soil moisture should increase reproductive investment (more flowers and larger flowers), but strong allocation trade-offs will produce a quantity-quality pattern in which plants with very high flower production may exhibit lower rewards (e.g.: nectar sugar concentration); because of resource dilution and risk of geitonogamy [31] (ii) Fruit set and conspecific pollen loads will be driven by individual visitation patterns; visitation will increase with rewards and flower size, with plants offering higher rewards attracting a more diverse visitor set, (iii) Increasing numbers of simultaneously flowering conspecifics will intensify competition for pollinators, reducing conspecific pollen deposition and fruit set at the focal plant (iv) The individual-based interaction structures are predicted to mirror microhabitat (soil moisture) gradients, where plants in wetter microsites that invest more in flowers and nectar will act as highly connected hubs with greater niche overlap; consequently, removing dominant pollinator groups will most strongly impact hub plants occupying favorable microhabitats, and targeted removal of highly connected individuals will disproportionately reduce generalization and overall robustness compared with random removals.

## Methods

### Study area

The study was conducted in the Aravalli ranges in Manethi, located in the Rewari district of Haryana, northwestern India (76.403739° E, 28.177195° N) (Fig. 2). The Aravalli ranges are among the oldest fold mountain systems in the world and are characterized by a semi-arid forest ecosystem. It acts as a boundary separating the desert in the west from the rich lowlands in the east. This region has an annual temperature range of 0-46°C and receives annual precipitation between 400-550 mm. There are around 300 native plant species, 120 bird species, and 398 insect species which are mainly comprised of Hymenoptera, Lepidoptera, and Coleoptera that were documented in the Aravallis [32]. The area has undergone a depletion of groundwater resources, driven by intensified anthropogenic activities in recent years [33].

**Fig. 2.**
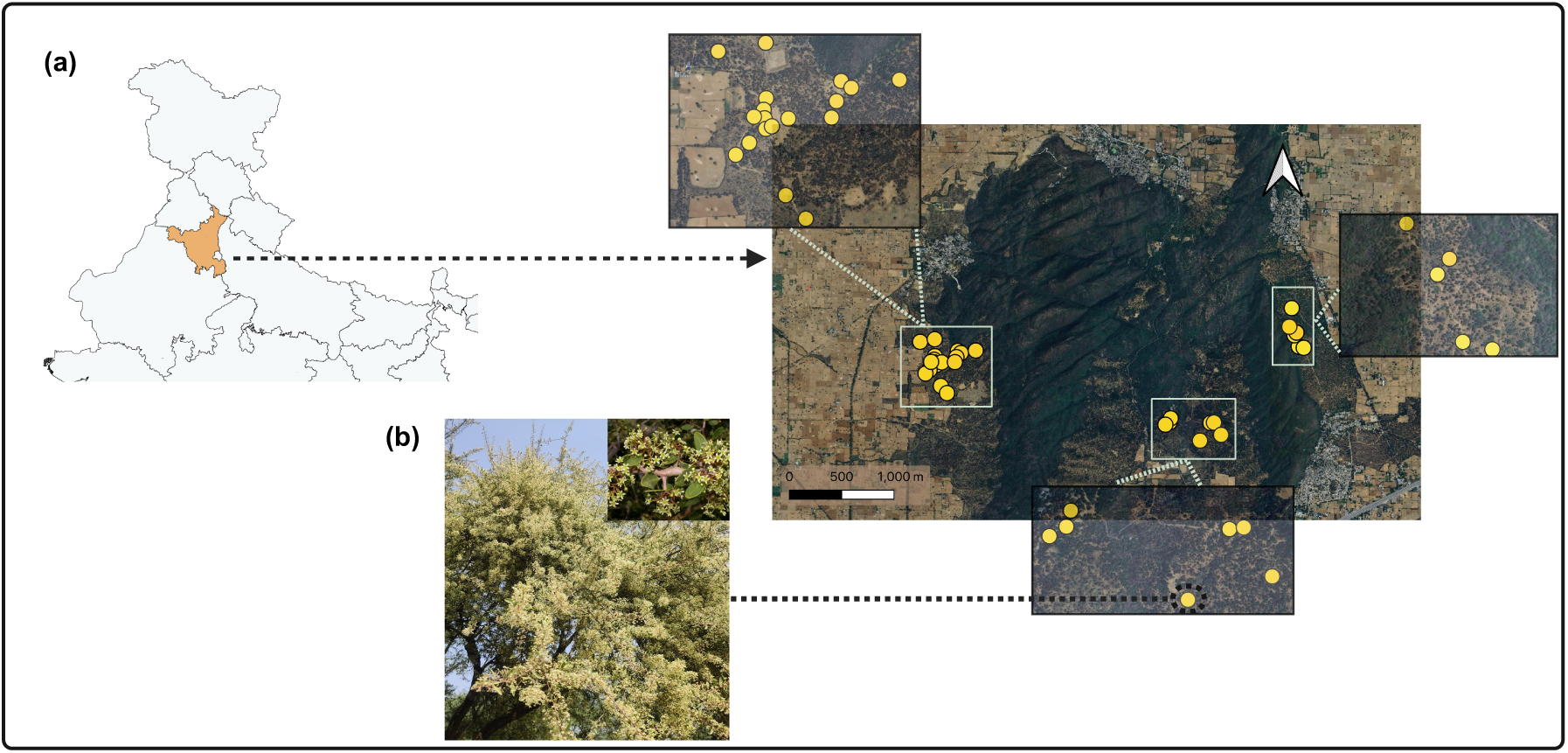
(a) Study location: Aravalli Hills, Manethi, Haryana, along with a satellite image of the Aravalli Hills in Manethi, where *Maytenus* individuals are marked as yellow dots according to their respective locations and insets showing the enlarged areas of the marked *Maytenus* plants (b) Image of a *Maytenus senegalensis* plant, with an inset showing the flower.

### Study species

The study species, *Maytenus senegalensis* (family Celastraceae), is a native and dominant shrub of the Aravalli ranges. It occurs across Africa, the Arabian Peninsula, Afghanistan, Pakistan, India, Bangladesh, and Spain. The plants are usually 2-6 meters in height and produce small, creamish-white flowers (Fig. 2). *M. senegalensis* relies largely on cross-pollination for reproductive success [34]. In our study area, the flowering period generally occurs between October and November, during which they exhibit a mass flowering pattern.

### Habitat factors

A total of 31 plants were selected based on the availability of the flowering individuals, as the species exhibits asynchronous flowering phenology among individuals within the community. Each selected plant was geotagged across the study site (Fig. 2). A minimum distance of 20-30 meters was maintained between individual plants. A 10 × 10 m plot was established around each focal plant (a schematic representing the plot is given in the additional file: section 1.1), and within this plot, the identity of neighboring plants and their phenological status were recorded to assess the influence of spatial neighborhood on the focal individual. Habitat factors, including soil moisture percentage and proximity to the edge of the forest, were measured (forest patch was surrounded by agricultural lands and roads as the forest boundaries) to assess habitat-level variations influencing plant traits.

Soil moisture percentage was measured using a soil moisture meter (Extech 750), a device that is equipped with a 20 cm probe that is inserted into the soil to provide moisture readings as percentage with measurement range from 0 to 50% (accuracy specified as ± 5% + 5 digits of full scale). Each month during the flowering period four moisture measurements were taken from different cardinal directions within a 2 m radius of each focal individual, and the mean was calculated. We used a 2 m radius to standardize the spatial scale of soil moisture close to the plants. The probe measures a very local volume, so taking four readings distributed around the plant within 2 m averages fine-scale heterogeneity in the microhabitat. The overall soil moisture percentage for each individual was calculated by averaging these monthly means measured from the flowering period to the fruit set stage (October to December 2023).

The distance to the habitat edge for each individual was calculated using QGIS software (version 3.34 Prizren) [35]. Proportion of conspecific neighbors were quantified as the proportion of flowering conspecific individuals within each 10 × 10 m plot created around each focal individual, where a value of 0 indicates the absence of co-flowered conspecifics and values approaching 1 indicate high proportion of conspecific neighbors.

### Floral resources

Floral traits and nectar rewards were measured for each focal individual. Seven floral morphometric traits, including petal length, petal width, flower diameter, flower height, stamen height, anther width, and stamen width, were measured using digital callipers with an accuracy of ±0.02 mm (n = 5 flowers/individual; 27 plants). These measurements were taken only once per plant and were conducted by a single observer. Flower production was estimated for all focal individuals by counting the number of flowers every 5-6 days using the quadrat method throughout the flowering period [36]. An imaginary quadrat of 0.5 × 0.5 m was created, and the number of flowers within it was counted (this count was then multiplied by the total number of possible quadrats that could fit within an individual plant’s area, based on its size). The total flower production per individual was calculated by summing these counts. Nectar sugar concentration was measured by bagging the buds to protect from the nectar robbery and pollinator visits. To minimize selection bias, flowers were selected at anthesis, ensuring they were undamaged and measured in % Brix using a handheld refractometer after removing the flowers from the plant (Cole-Parmer® Brix Analog Refractometer; n = 8-12 flowers/individual; 28 plants).

### Plant-pollinator observations

Flower visitation observations were conducted in 15-minute intervals during the flowering period on all the focal individuals (30-120 minutes/individual over 34 days). In each 15-minute slot, the number of pollinators, the number of pollinator visits, and the identity of the pollinators were recorded. Observations were conducted during both morning (7:00 AM–1:00 PM) and evening (3:30 PM–6:30 PM) hours by a single observer. A total of 31 plant individuals were sampled. Each individual was monitored multiple times throughout the flowering period based on its flower production. A visit was defined as a pollinator making contact with the reproductive parts of a flower. The majority of pollinators were identified to the genus level. The abundance of *Apis* pollinators was 9.07%, while *Eristalinus* pollinators comprised 17.9% of the total pollinators observed. Moreover, the genera *Apis* (Hymenoptera) and *Eristalinus* (Diptera) accounted for 35.36% and 22.45% of the total pollinator visitations, respectively. Pollinators were classified into 78 morphotypes, of which 48 were identified to the genus level, 7 to the family level, and 23 to the order level.

### Assessment of fruit set and pollen loads

The reproductive success of the plants was assessed by measuring conspecific pollen loads on flower stigmas (proxy for pollination success) and calculating the percentage fruit set. To quantify the conspecific pollen loads, flowers were collected immediately after pollinator visitation observations (15 min slots), and only flowers visited by pollinators were selected (n = 25 flowers/individual; 18 plants). Stigma collections were carried out by the same two observers who recorded the pollinator visitations. For each individual, 25 stigmas were collected across multiple observation days throughout the flowering period to avoid any bias. The stigma from each flower was removed and visualized using the stigma squash method with basic-fuchsin jelly [37]. To distinguish conspecific pollen from heterospecific pollen, reference pollen grains from the anthers of *M. senegalensis* were observed before analysis.

The percentage fruit set was determined by tagging mature flower buds (n = 10-15 buds/individual; 31 plants) based on their availability. All the selected buds on each individual were marked on the same day and monitored their development every 5-6 days to record their transition from buds to flowers and subsequently to immature fruits. For each plant, this was calculated as the ratio of immature fruits formed to the total number of tagged flowers.

## Analysis

All statistical analyses and plot visualizations were performed using R software (version 4.4.2) [38] in RStudio (version 2024.09.1+394) [39]. The relationships among traits such as flower diameter, petal width, petal length, flower height, stamen height, anther width, and stigma width were analysed using principal component analysis (PCA). Among these traits, flower diameter and petal width had the highest principal component (PC) scores. Given the correlation between flower diameter and petal width (Pearson correlation: cor = 0.78, *p* < 0.001), we included only flower diameter in further analyses. A summary of sample sizes and variables used in the analyses is given in supplementary methods (Additional file: section 1.3).

We analyzed the correlation between floral traits and soil moisture to assess whether variation in soil moisture affects the floral traits of individual plants, and also examined the correlations among these traits to investigate potential trait trade-offs. We assessed the spatial distribution of individual plants using Moran’s I to test for spatial autocorrelation, using the spdep [40] and sf packages [41] in R. We analyzed the sampling completeness of pollinator abundance, genera richness, and pollinator visitation using the iNEXT function, evaluate sufficiency of sampling effort [42]. To investigate variation in pollinator interactions among *Maytenus* individuals, we constructed an individual-level interaction network. In this network, plant individuals of *Maytenus* were considered as nodes and connected to pollinators, and the number of visits was considered for the connection weights. Network-level and node-level metrics were calculated using the “networklevel” and “specieslevel” functions [43, 44]. All the analyses related to networks were conducted using the bipartite package [45].

We evaluated whether the observed network structure is a result of random processes or exhibits underlying patterns influenced by ecological factors, such as habitat or the availability of floral resources. For this, we employed the vaznull algorithm to generate 999 null models with constrained connectance and moderately constrained marginal totals [46]. Network-level metrics were extracted from these null models and compared to the metrics derived from the observed network. To quantify the deviation of the observed network metrics from those of the null models, *z*-scores were calculated, providing a measure of the variation between the observed and randomized networks. The extracted network metrics include weighted connectance, Shannon diversity, niche overlap, linkage density, weighted NODF, and H2. A brief overview of these metrics is below:

a. Weighted connectance quantifies the proportion of realized interactions in a network. The values range from 0 (no realized interactions) to 1 (all possible interactions fully realized with maximum weights).
b. Shannon diversity measures interaction diversity by incorporating both richness (number of interactions) and evenness (distribution of interaction strengths).
c. Linkage density represents the average number of interactions per plant individual, weighted by interaction strengths, indicating network connectivity; values greater than 0 suggest denser, more connected networks.
d. H2 quantifies the degree of specialization in the network, with values spanning from 0 (completely generalized network with random interactions) to 1 (highly specialized network with unique, specific interactions).
e. Weighted NODF extends the understanding of nestedness in networks to account for interaction weights, with values ranging from 0 (random, unorganized interactions) to 100 (strongly nested networks where interaction presence and strength align with nested patterns).

Analyzing the modularity of a network allows us to determine whether the network is divided into distinct groups or modules, within which plants and pollinators interact more strongly with each other than with those in other modules. To analyze the modularity of the network, we utilized the Dormann Strauss algorithm in “metaComputeModules” function (99 replications) [47]. To understand the variation between the modules in reproductive success, the Kruskal-Wallis test, followed by Dunn’s pairwise comparisons test, was used. The stability of the network against perturbations was assessed by removing the dominant pollinator groups and comparing those network metrics with the full networks and by removal of individual plants [48]. Removal of plants was done using the “second.extinct” function, which simulates species loss under two scenarios: (1) individual plants were removed in a random order, and (2) individual plants were sequentially removed based on their degree. We calculated the exponent of the hyperbolic function curve fitted to the proportion of secondary extinctions when individual plants were removed.

Paired Differences Index (PDI), normalized degree, species strength, weighted closeness, and d were computed to assess interaction differences between individual plants. A brief overview of these metrics is below:-

a. PDI quantifies the degree of specialization of an individual within a network by assessing how its interaction strengths differ from those with other potential partners. It emphasizes asymmetries in interaction strengths, with values ranging from 0 to 1. A value of 0 indicates complete generalization, where interaction strengths are evenly distributed across partners, whereas a value of 1 reflects complete specialization, where a species strongly interacts with only one partner.
b. Normalized degree represents the degree of a plant as a proportion of the total possible degree, scaling the plant’s connectivity relative to all potential connections. Values range from 0 (no connections) to 1 (connected to all other nodes in the network).
c. Species strength measures the total interaction strength of a plant individual by summing the weights of its interactions with all partners. This metric reflects the individual plant’s importance in the network, with values starting at 0 (indicating no interactions). Higher values correspond to greater total interaction strength.
d. Weighted closeness centrality measures the efficiency with which a plant can connect to other plants in the network through shared pollinators, considering both the number of connections and the interaction strengths. Plants with higher weighted closeness centrality are more centrally positioned in the network, indicating stronger and more connections to other plants via shared pollinators. Lower values suggest fewer shared pollinator interactions between the plants.

Apart from these node-level metrics, we also calculated the interaction niche overlap between plant pairs within the network (niche.overlap function, Schoener’s index, spaa package) [49]. A binary incidence matrix with the presence and absence of pollinator genera was used for the calculation of niche overlap values. Niche overlap provides insight into the extent to which a plant shares pollinator resources with other plants in the network. Niche overlap values for each plant individual were then analyzed using a one-sample Wilcoxon test to assess whether any individual’s niche overlap significantly deviates from the median value across all individuals.

In our study, binary Exponential Random Graph Models (ERGMs) were used to investigate how various predictors influence the presence and absence of pollinator visits to plant individuals (ergm package) [50]. ERGMs are similar to Generalized Linear Models (GLMs) but are specifically designed for network data. Binary ERGMs employ maximum pseudolikelihood estimation (MPLE) to estimate the effects of predictor variables on the likelihood of connections (edges) in the network. All predictor variables in this analysis were attributed as node-level covariates.

Multiple GLMs were constructed with percentage fruit set as response variable. Flower diameter, nectar sugar concentration, flower production, soil moisture, distance to habitat edge, and proportion of conspecific neighbors were considered as predictors (details of model construction and model diagnostics are outlined under the statistical analysis section of the additional file). Apart from these variables, we also constructed models with pollinator abundance and number of pollinator visits as predictors.

Generalized Linear Mixed Models (GLMMs) were used to assess the effect of floral traits and habitat factors on conspecific pollen load receipt. Since variation within plants could contribute to some of the observed effects, we used plant identity as a random effect for the analysis. Flower diameter, nectar sugar concentration, flower production, soil moisture, distance to habitat edge, and proportion of conspecific neighbors were considered as predictors. GLMs with pollinator abundance and number of visits as predictors were fitted, as models without a random effect provided considerably better fit (details in the Additional file).

To ensure appropriate model fit, variables were log-transformed where necessary. The model selection process was guided by the Akaike Information Criterion (AIC), with the model that had the lowest AIC being considered the best fit. Additionally, variables were iteratively reduced based on their significance levels, determined through ANOVA (car package) [51]. These models were fit using the glmmTMB() function in the glmmTMB package [52]. The goodness-of-fit for both GLM and GLMM was evaluated using the simulateResiduals() function from the DHARMa package [53].

## Results

### Intraspecific variation in floral resources

The first two principal components (PC1 and PC2) accounted for the majority of the variation in the floral traits, explaining 54.8% and 14.09% of the total variance, respectively. The estimated flower production exhibited substantial variation among plant individuals, ranging from 83,000 to 1,100,000 flowers (coefficient of variation, CV = 63.57%). Nectar sugar concentration levels varied between 3.0% and 6.75% Brix across individuals (CV = 20.47%), while flower diameter ranged from 3.4 to 5.82 mm (CV = 15.75%). CV values indicate that flower production exhibited the highest variability among the traits (Fig. 3a). We observed that flower diameter increased significantly with soil moisture. Flower production also trended upward, whereas nectar concentration showed no significant association with moisture. (Additional file: Fig. S1, Table S1).

**Fig. 3.**
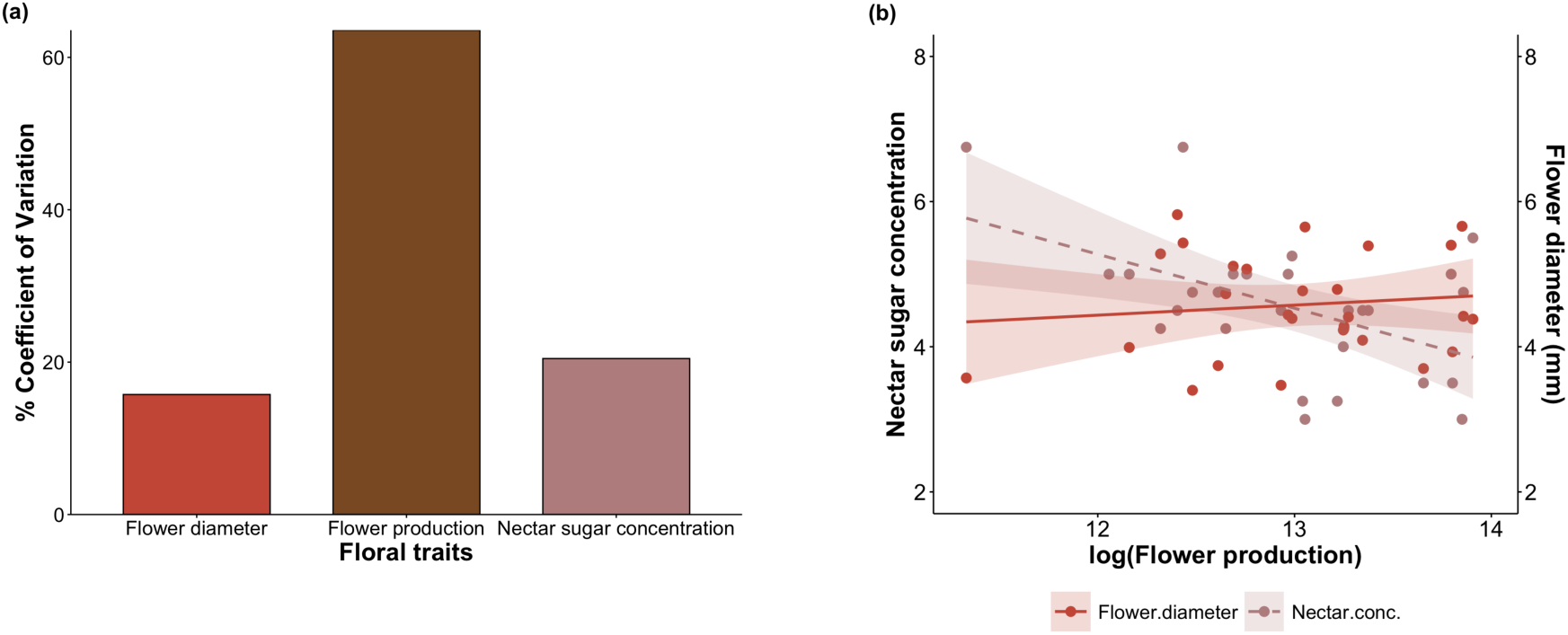
(a) Bar plot depicting the variation in floral traits among *Maytenus* individuals, expressed as the percentage coefficient of variation (% CV) for each trait. (b) Scatter plots depicting the variability in median nectar sugar concentration (% Brix) and mean flower diameter (mm) with increasing flower production in *M. senegalensis* individuals. Pearson correlation was used to assess the relationships between the floral traits. Each dot represents an individual plant, and the dashed line indicates the significant relationship.

To investigate potential trade-offs in resource investment, we examined the relationships between flower production, flower diameter, and nectar sugar concentration. We observed a negative correlation between flower production and nectar concentration (Pearson correlation: cor = −0.49, *p* = 0.007). On the other hand, flower diameter was not significantly related to changes in nectar concentration or flower production (Fig. 3b, Additional file: Table S2).

### Plant-pollinator interaction structures

Sampling completeness values for the abundance of pollinators and the interactions were found to be greater than 98%, indicating sufficient sampling of the individuals (Additional file: Fig. S2). A total of 7,458 pollinator visits were recorded for the focal plants, with visits largely from the pollinator orders, Diptera and Hymenoptera. The most dominant visitors were from the genera *Apis* (Hymenoptera) and *Eristalinus* (Diptera), accounting for 35.36% and 22.45% of the total recorded visitation, respectively (Fig. 4). When comparing the observed networks to random assemblages, all the null model metrics differed considerably from the observed network (Additional file: Fig. S3, Table S3). All network metrics, except for H2, exhibited negative *z*-scores, indicating that their observed values were lower than the predicted averages. In contrast, H2 displayed a positive *z*-score, suggesting that its observed value exceeded the expected value from the null models. This non-random assemblage pattern was further analysed using ERGMs to examine the attributes that contribute to it. Binary ERGMs suggest that flower production, distance to the habitat edge, and flower diameter negatively influenced the presence of a connection with a pollinator genera, while soil moisture had a positive effect. Nectar sugar concentration and proportion of conspecific neighbors were not found to have an effect (Table 1).

**Fig. 4.**
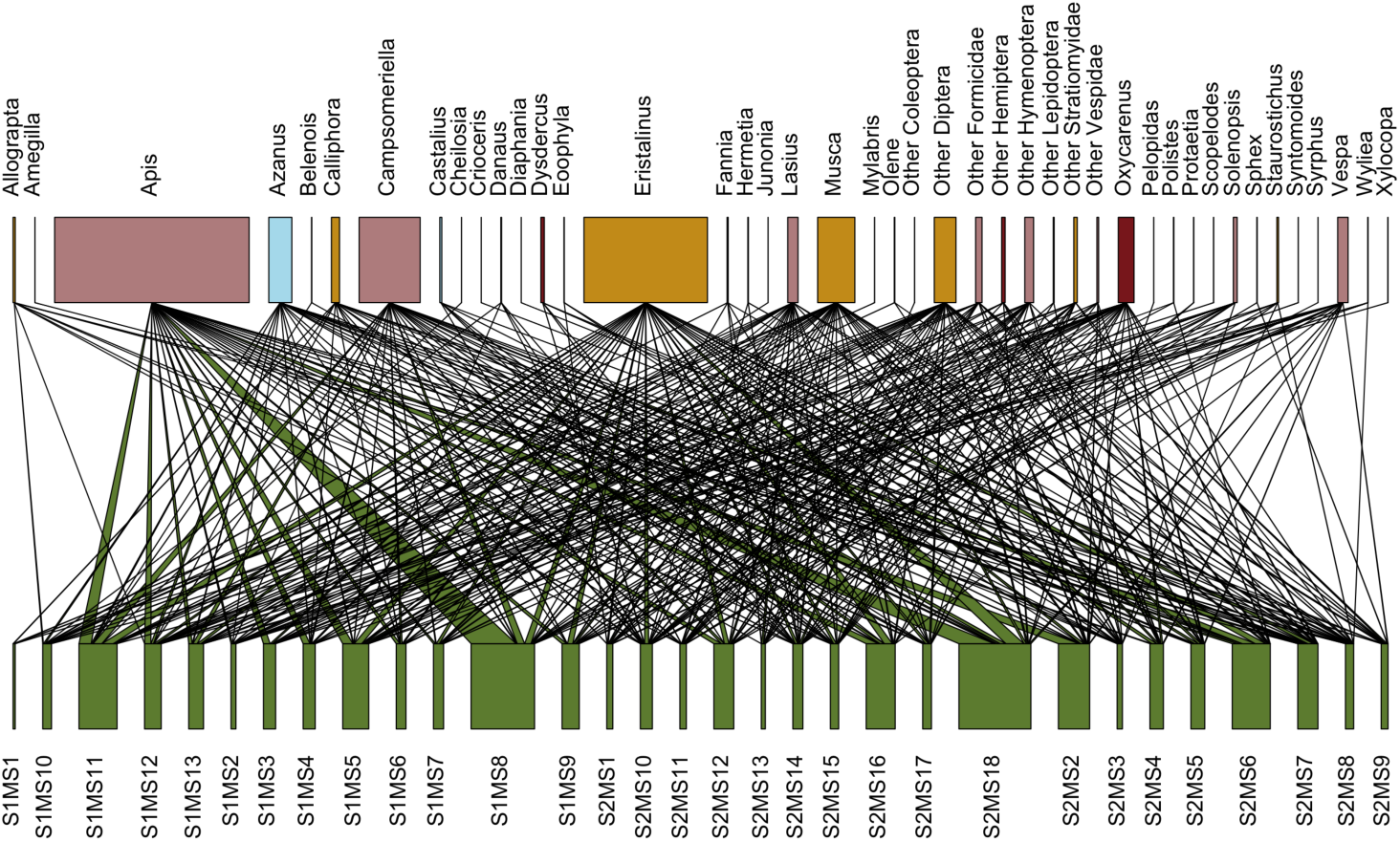
Weighted bipartite network showing the interactions between pollinator species and *Maytenus senegalensis* individuals (green). Pollinator species are colored according to their taxonomic order as follows: Hymenoptera – rosy brown; Diptera – yellow; Lepidoptera – blue; Coleoptera – gray; Hemiptera – brown. The links between nodes represent flower visitation interactions, with the width of the links corresponding to the number of visits recorded. The pollinators mentioned as ‘Other’ are the morphotypes that could not be identified until the genus level (e.g., Other Coleoptera).

**Table 1.** Summary of the binary exponential random graph models (ERGM) showing the effects of individual plant floral traits and habitat factors on the structure of the bipartite plant-pollinator network. Estimates of predictor variables with significant effects indicate the expected change in the presence of plant-pollinator interactions given a one-unit change in that predictor variable. Significant values (*p* < 0.05) appear in boldface.

|  | Predictor variables | Estimate | Std.Error | z value | p-value |
| --- | --- | --- | --- | --- | --- |
| Edges | Nectar concentration | 0.01 | 0.04 | 0.39 | 0.68 |
|  | Flower diameter | -0.14 | 0.04 | -3.04 | **0.002** |
|  | Soil moisture | 0.29 | 0.05 | 5.62 | **<0.001** |
|  | Flower production | -0.22 | 0.05 | -4.27 | **<0.001** |
|  | Proportion of conspecific neighbors | 0.03 | 0.04 | 0.69 | 0.48 |
|  | Distance to habitat edge | -0.24 | 0.04 | -4.96 | **<0.001** |

To evaluate the impact of the dominant insect orders on network structure, we also constructed networks excluding each of them individually (Additional file: Fig. S4, Fig. S5, Table S4). The specialization index (H2), weighted NODF, and linkage density exhibited substantial variation between the full network and the network without Diptera, with differences exceeding 20%. The removal of Diptera increased specialization and lowered linkage density and weighted NODF. In contrast, these metrics remained comparable between the full network and the network without Hymenoptera. Weighted connectance was 25% higher in the absence of Hymenoptera compared to the full network. The PDI values indicated that individual plants exhibited a high degree of specialization, with values ranging between 0.92 and 0.98. The mean normalized degree value was 0.23. Species strength range varied from 0.12 to 3.28. Five individual plants were identified as key contributors to the network’s structure (values higher than 2). The mean weighted closeness centrality value of plants was 0.03. The niche overlap values of plants differed significantly, with a few individuals exhibiting low niche overlap (< 45%), indicating minimal sharing of pollinators (One-sample Wilcoxon test: V= 59, *p* = 0.02, Fig. 5; Additional file: Table S5). We assessed the correlations between species strength and floral traits, as well as niche overlap and floral traits, and found no relationship (Additional file: Fig. S6, Table S6).

**Fig. 5.**
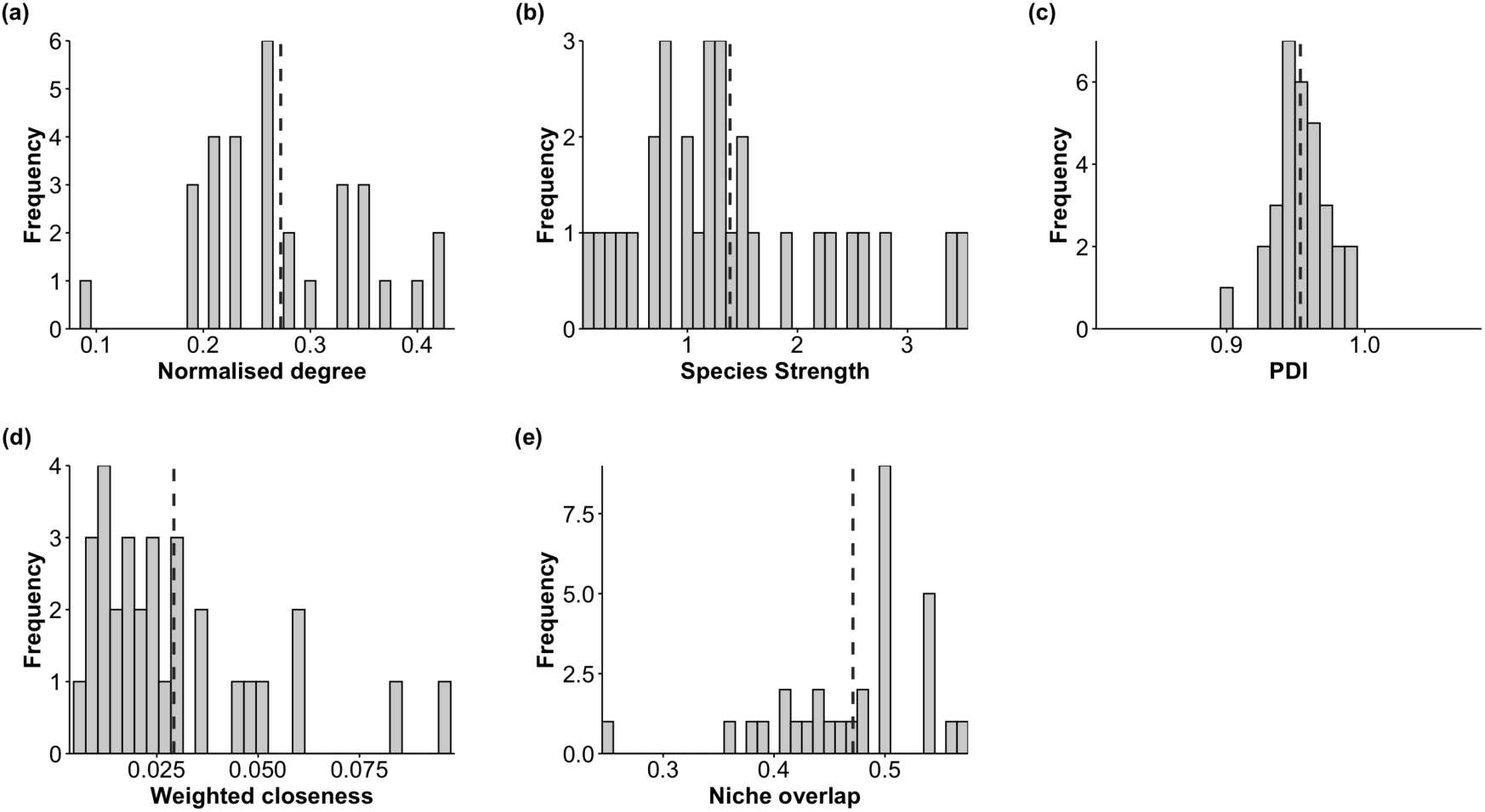
Bar graphs depicting the plant-level metrics from the network. Each bar represents the frequency of plants with the corresponding metric, with the dashed line indicating the mean value of the metric. Network metrics include: (a) Normalized degree, (b) Species strength, (c) Paired Differences Index (PDI), (d) Weighted closeness, and (e) Niche overlap (Schoener’s index).

The analysis of the observed network revealed a low modularity value of 0.227, with one large module containing most plant individuals and two smaller modules (Additional file: Fig. S7a, Fig. S7b). The robustness values were 0.737 for random removal and 0.612 for degree-based removal of individual plants. The exponent values were 3.72 for random removal and 1.83 for degree-based removal. Overall, the results show that the network is moderately resilient to individual plant removal, and the removal of plants with the highest interactions might influence the stability of the network (Additional file: Fig. S7c, Fig. S7d).

### Intraspecific variation in reproductive success

The median conspecific pollen loads across individuals ranged from 13 to 431 (SD = 99.21, Mean = 95.08). The standard deviation of intra-individual pollen loads exhibited considerable variability, with certain individuals displaying minimal variation in pollen load while others showed greater intra-individual variation. Generalized Linear Mixed Model (GLMM) analysis indicated that flower diameter and soil moisture had a positive effect on conspecific pollen loads. Flower production also showed a marginal positive effect on pollen loads, whereas, proportion of conspecific neighbors had a negative effect on conspecific pollen loads (Fig. 6, Table 2). The nectar concentration and distance to habitat edge did not affect the conspecific pollen loads within individual plants. (Fig. 6, Table 2). To further examine the role of inter-individual variation, a GLM was constructed using the median conspecific pollen loads per individual, and the analysis showed a similar trend (Additional file: Table S7). Visitation rate and pollinator abundance did not show any significant effect on median conspecific pollen loads per individual (Additional file: Fig. S8, Table S8).

**Fig. 6.**
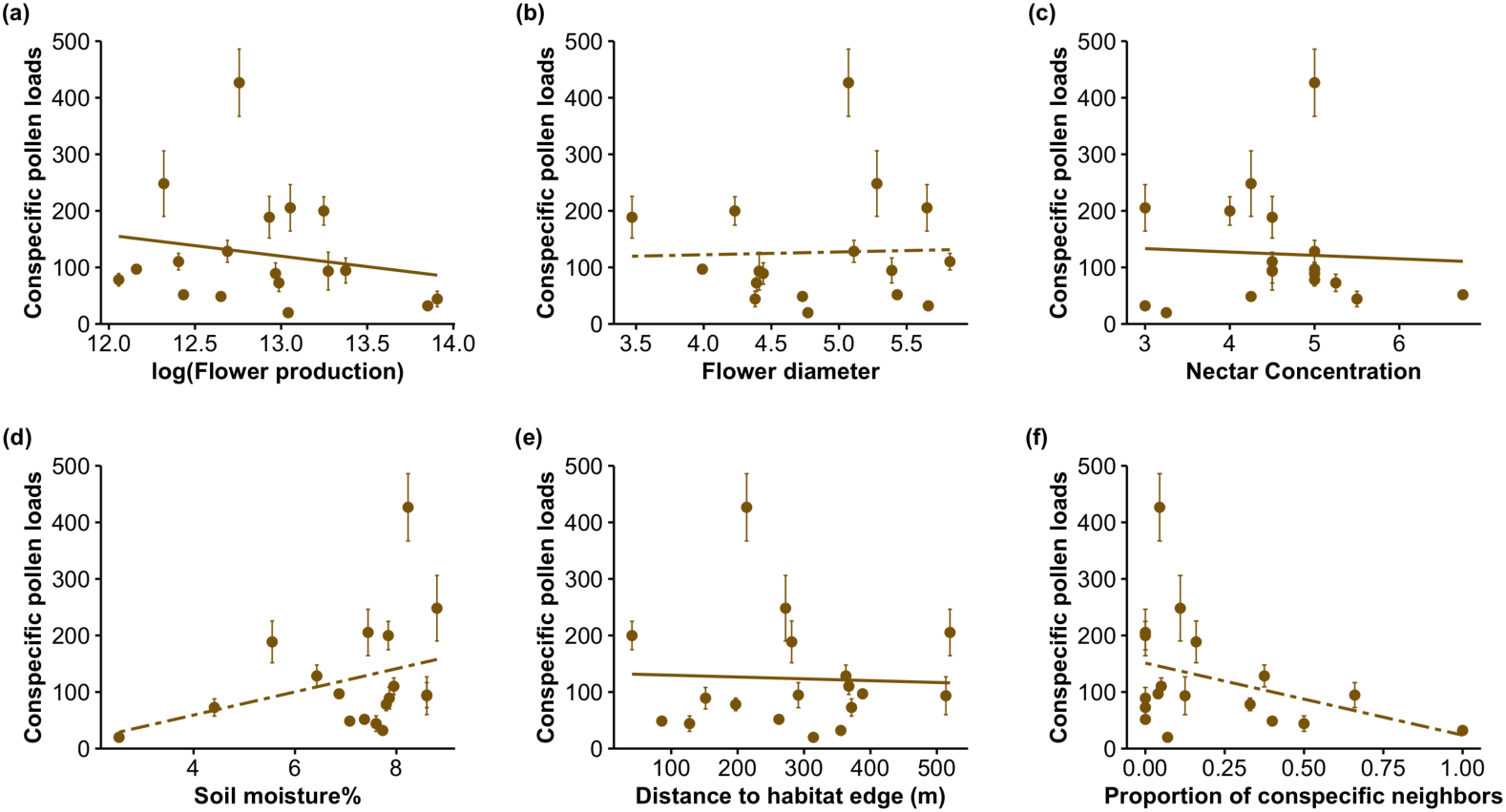
Scatter plots illustrating the variability in conspecific pollen loads of individual plants, explained by (a) flower production, (b) flower diameter, (c) nectar sugar concentration (% Brix), (d) soil moisture %, (e) distance to habitat edge, (f) proportion of conspecific neighbors. Each point represents the mean and ± SE of an individual plant, and the dashed fit line indicates the significant relationship.

**Table 2.** GLM of percentage fruit set and GLMM of conspecific pollen loads in response to floral traits (flower production, nectar sugar concentration, flower diameter) and habitat factors (soil moisture %, proportion of conspecific neighbors and distance to habitat edge) of *M. senegalensis* plants. Flower diameter and nectar concentration were measured on multiple flowers per individual plant; for the GLM, plant-level predictors were the mean flower diameter and the median nectar concentration. Both response variables were log-transformed, and Gaussian family was used for all models. Predictor variables that were scaled are indicated by (*z*) in the table. Significant values (*p* < 0.05) appear in boldface.

| Response variables | Models | Predictors | Estimate | SE | *z* | *p* | Random effect | SD (RE) | Var. (RE) |
| --- | --- | --- | --- | --- | --- | --- | --- | --- | --- |
| Fruit set% | Model 1 | Nectar Concentration | -0.088 | 0.268 | -0.328 | 0.742 |  |  |  |
|  |  | Flower diameter | -0.316 | 0.306 | -1.030 | 0.303 |  |  |  |
|  |  | log(Flower production) | -1.291 | 0.410 | -3.149 | **0.001** |  |  |  |
|  | Model 2 | Soil moisture | -0.307 | 0.149 | -2.054 | **0.039** |  |  |  |
|  |  | Distance to habitat edge | 0.002 | 0.001 | 1.435 | 0.151 |  |  |  |
|  |  | Proportion of conspecific neighbors | -1.294 | 0.985 | -1.314 | 0.188 |  |  |  |
| Conspecific pollen loads | Model 3 | Soil moisture | 0.251 | 0.089 | 2.814 | **0.004** |  |  |  |
| | | Distance to habitat edge | $0.8 \times 10^{-4}$ | 0.001 | 0.084 | 0.932 | Plant ID | 0.537 | 0.288 |
|  |  | Proportion of conspecific neighbors | -1.469 | 0.496 | -2.961 | **0.003** |  |  |  |
|  | Model 4 | Nectar Concentration (*z*) | -0.206 | 0.143 | -1.442 | 0.149 |  |  |  |
|  |  | Flower diameter (*z*) | 0.314 | 0.102 | 3.075 | **0.002** | Plant ID | 0.547 | 0.3 |
|  |  | log(Flower production) | -0.634 | 0.343 | -1.845 | 0.064 |  |  |  |

Fruit set percentages ranged from 0% to 100% (mean ± SD = 22.25 ± 33.24%), with most plants exhibiting low fruit sets. GLM analysis showed that the flower production and soil moisture have a significant negative influence on the percentage fruit set (Fig. 7, Table 2). Visitation rate and, marginally, pollinator abundance were positively associated with fruit set, but neither significantly explained conspecific pollen loads (Additional file: Fig. S8, Table S8). There was considerable variation in the percentage of fruit set between the network modules (χ² = 33.81, df = 2, *p* < 0.001). Post-hoc Dunn’s tests revealed differences in reproductive success between Module 1 and Module 2 (*p* < 0.001) and between Module 2 and Module 3 (*p* < 0.001). However, no differences were found between Module 1 and Module 3 (*p* = 0.33).

**Fig. 7.**
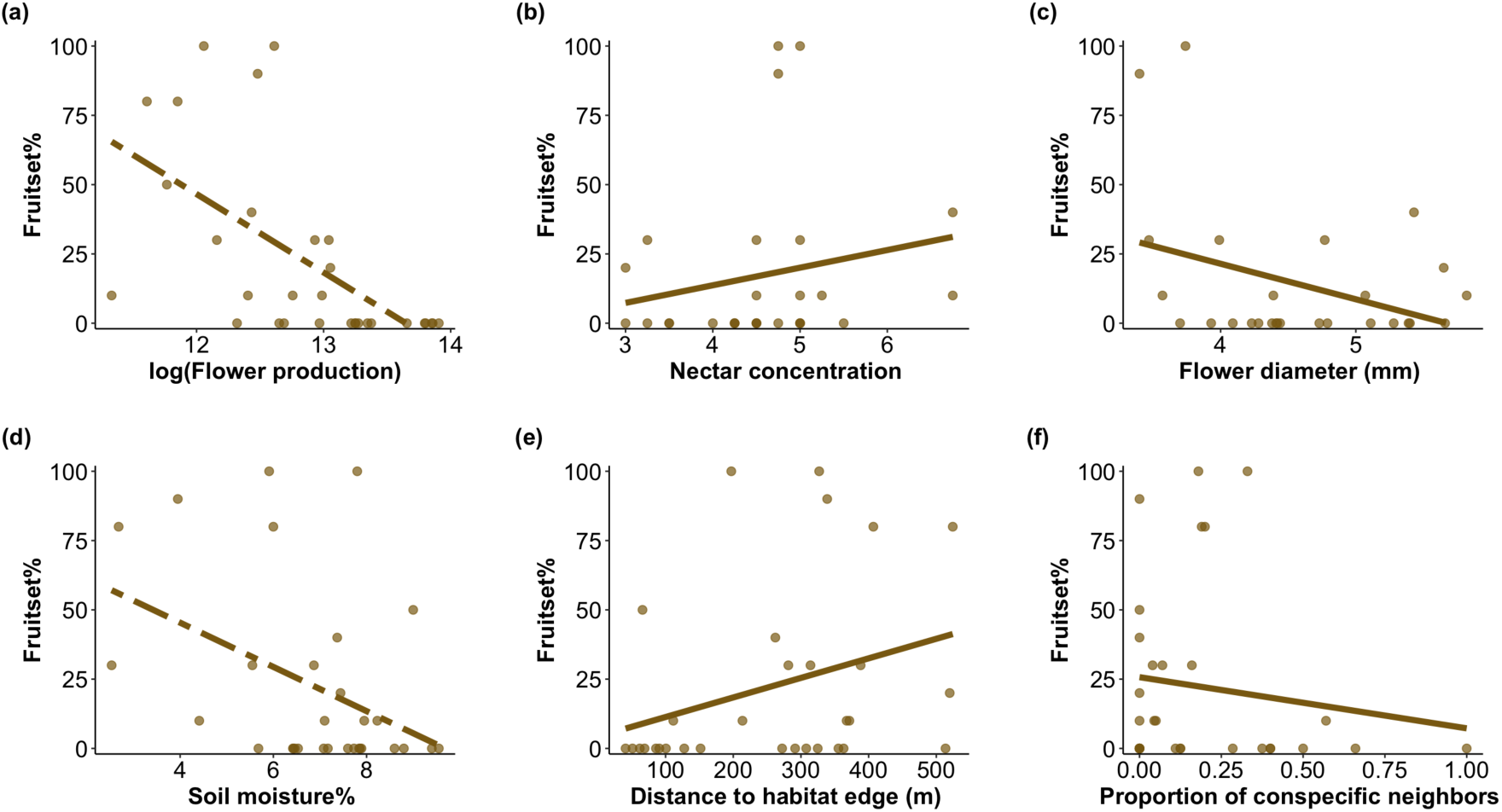
Scatter plots illustrating the variability in fruit set % of individual plants, explained by (a) flower production, (b) nectar sugar concentration (% Brix), (c) flower diameter, (d) soil moisture %, (e) distance to habitat edge, (f) proportion of conspecific neighbors. Each point represents an individual plant, and the dashed fit line indicates the significant relationship.

## Discussion

Our study investigated the relationships between floral traits, habitat factors, and plant-pollinator interactions in a semi-arid region, revealing substantial inter-individual variation in flower production and a trade-off with nectar sugar concentration. Soil moisture was positively associated with flower diameter (and, to an extent, flower production) but was negatively associated with fruit set. Flower production negatively influenced pollinator interactions, thereby reducing reproductive success. Pollinator interaction structures were further shaped by flower diameter, soil moisture, and neighborhood composition: flower diameter and soil moisture increased conspecific pollen loads, whereas a higher proportion of conspecific neighbors decreased them. Additionally, while individual plants occupied overlapping pollinator niches, network analyses indicated a degree of specialization likely driven by competition avoidance. Furthermore, stability analyses showed that Dipterans were critical for network structure, and simulated plant removals revealed that the loss of highly connected plant individuals had a greater impact on individual-level network resilience.

Resource allocation patterns are largely genetically determined within plant species but can be influenced by environmental conditions [54]. Specifically, plants in resource-limited environments were often found to exhibit selective resource allocation strategies that enhance reproductive success [55, 56]. However, the extent to which microhabitat variations impact plants and their flower visitors was variable [13, 19, 21]. In this study, conducted in a semi-arid region, we hypothesized that *Maytenus senegalensis* individuals would exhibit a reduction in floral traits with decreasing soil moisture levels. We found that soil moisture variation affected flower size and production, and there was considerable intraspecific variation in floral traits. Individuals with higher flower production tended to have lower nectar sugar concentration, while those with fewer flowers exhibited higher nectar sugar concentration. Flower production varied negatively with nectar sugar concentration, suggesting a trade-off in the allocation of resources between these traits [57, 58]. This could be expected when the energetic costs involved in the production of nectar sugars are higher than the costs of producing flowers [59, 60]. Such trade-offs that are indicative of quality (reward) and quantity (number of flowers) allow the maintenance of the combined attraction of the plant for the pollinators [61, 62]. Overall, we also found that there was higher variation at the individual level in flower production, suggesting higher plasticity in this trait [63, 64].

Pollinators that are abundant in the local habitat and visit multiple plants are known to be important for the stability of the network [65]. In our study, the exclusion of pollinators from Diptera resulted in a more specialized network with lower nestedness values and linkage density, indicating a reduction in generalization and connectivity. Such a pattern suggests that Dipterans are critical for *M. senegalensis* in maintaining network generalization. On the other hand, weighted connectance exhibited significant increases following the removal of Hymenoptera and was relatively unaffected by the removal of Diptera. In a network without Hymenopterans, pollinator interactions were found to increase, this could potentially be driven by reduced interspecific competition between pollinators and niche displacement by other pollinator groups. These findings indicate that the functional role of Hymenoptera within the network can be partially compensated by other pollinators. Consistent with our prediction (*iv*), the strongest reductions in generalization following Dipteran removal were concentrated on highly connected “hub” plants, particularly those occupying wetter microsites that supported larger displays and higher nectar rewards.

Our observed network differed from the randomly generated networks and was found to exhibit a higher degree of specialization. The higher specialization is likely driven by ecological factors such as pollinator preferences and resource partitioning [66]. ERGM analyses suggest that traits such as flower production and flower size are critical in determining the likelihood of a particular type of pollinator visiting the plant. A couple of broad patterns in pollinator visitation were seen in our data. Plants visited by pollinators from multiple genera were found to have flower production or nectar sugar concentration that is slightly higher than others, whereas plants visited by pollinators from fewer genera showed intermediate levels of both traits. Therefore, as anticipated in prediction *(ii)*, visitation tended to respond to nectar sugar concentration and flower diameter, with higher-reward plants attracting a more diverse visitor set, thereby increasing the potential for effective pollen transfer. These observations suggest a possible relationship between genera richness and floral traits, which requires further consideration.

In our study, individual plants were found to exhibit a high degree of specialization (high PDI values). The analysis showed that not all plants were connected within the network through shared pollinators. Some plants exhibited higher species strength values, and these individuals were found to have slightly higher nectar sugar concentrations, low flower diameter, and flower production than other plants. However, these trends are only indicative, as there were no statistically significant relationships with floral traits. Ten plant individuals within the network were found to have a low niche overlap (<45% overlap), pointing towards resource partitioning, possibly reducing competition between the plants [67]. The exponent values from the random simulation scenario indicate that the extent of redundancy in pollinator interactions was higher during initial removals, hence, the network remained stable, which was then followed by an abrupt collapse. On the contrary, the lower exponent values of degree-based removal suggest a collapsing trend from the first few removals itself. These patterns are similar to previous findings, where the loss of highly connected plants accelerated network collapse [15, 20]. This individual-level specialization and sensitivity to targeted removals align with prediction *(iv)* and possibly suggest that pollinator-mediated competition can drive divergence of interaction niches across individuals (prediction *ii*). Together, these results highlight the interconnectedness between individual-level variation in reproductive traits and network-level stability. Conserving highly connected plants is essential not only for maintaining network robustness but also for preserving the ecological processes that support reproductive success within the system.

The floral traits of plants are known to influence the modularity of plant-pollinator networks at the species level [68, 69]; however, this may not always hold at the individual level. The network in our study resulted in three modules. Within these modules, plants in one of the smaller modules (module 3) were found to be from higher soil moisture levels and were visited by Dipterans. This may be linked to higher flower production in plants found in areas with elevated soil moisture. However, previous studies reported that Dipterans preferentially selected flowers based on size or nectar availability rather than flower production [22, 23, 24]. Plants in the other smaller module (module 2) were surrounded by fewer flowering conspecifics, and their visits were largely from Hymenopterans. This pattern could be due to the change in foraging preferences of pollinators with the change in spatial distribution of floral resources and also to possibly avoid competition [70, 71]. Previous studies have demonstrated that modules dominated by abundant and common species tend to exhibit higher reproductive success [20, 72]. Our study revealed variation in reproductive success among the modules, where one of the smaller modules (module 2) had a significantly higher percentage of fruit set compared to the other two modules in the network. These module-level patterns reflect the microhabitat gradient (prediction *iv*), where wetter-microsite plants functioned as interaction hubs with greater niche overlap and stronger dependence on Dipterans, whereas lowconspecific-density plants were more closely associated with Hymenopterans.

A higher proportion of conspecific neighbors and greater pollinator abundance were both associated with lower conspecific pollen loads at focal plants. This finding supports our hypothesis that an increase in conspecific density intensifies competition among individuals. Soil moisture and flower diameter both had a positive effect on conspecific pollen loads. The trend of increasing flower size under higher soil moisture conditions may likely release and receive more pollen loads [73]. Partly in line with prediction (*ii*), variation in visitation helped mediate these links from traits to conspecific pollen loads and ultimately to fruit set. However, other floral traits and distance to habitat edge did not influence the conspecific pollen loads considerably. This lack of effect may be due to pollinator saturation or stigma saturation.

Despite the positive effects of moisture and flower size on pollen loads, fruit set declined with higher flower production and higher soil moisture, patterns consistent with a quantity–quality trade-off (prediction *i*) in which high flowering intensity reduces per-flower reward quality (e.g., lower nectar sugar concentration) and depresses effective pollination [74, 75]. Similar dynamics have been reported in the dry grasslands of Gansu Province, China, where a study on *Medicago sativa* found that pollen limitation was the major reason for the lowered reproductive success of the plant [74]. A study on *Senita cacti* in Arizona showed that higher flower production lowered the pollinators-to-flower ratio and caused fruit abortion [76]. We predict that this is linked to the observed resource trade-off between flower production and nectar concentration. Together, these results link microhabitat to interaction outcomes, where plants in wetter microsites tend to produce larger flowers and attract visitors, yet high local conspecific density and high flower production can reduce conspecific pollen deposition and fruit set; patterns that also help explain why well-watered, highly connected plants may function as hubs in the network while remaining vulnerable to competition and resource dilution (prediction *iv*).

Our study examined how microhabitat conditions and floral traits shape pollinator visitation patterns and reproduction in a keystone semi-arid shrub species. Because inference is based on a single focal species and season, we are cautious of widespread generalizations. Robust estimates of individual-level variations in pollinator interactions at the plant scale will require applying this framework across multiple species and years, while incorporating additional microhabitat axes. A few of the methodological caveats include trait measurements taken at different times of day, which may confound diurnal light and temperature effects, and variable sample sizes due to the short flowering period, which may reduce power and bias effect estimates despite our use of statistical methods tolerant to missing values. Soil nutrients, also drivers of floral traits, were beyond the scope of our study. Integrated measurements of nutrients and moisture would strengthen the findings. Given the primacy of water in semi-arid systems, we focused on soil moisture. Adding other factors such as nutrient availability, canopy cover, and proximity to edges, and tracking seasonal shifts in pollinator niches, will better resolve temporal dynamics and deepen our understanding of the pathways linking microhabitat to reproductive success.

## Conclusions

Our study shows the intricate links between floral traits, habitat factors, and plant-pollinator interactions in shaping the structure and resilience of ecological networks. Key findings include the critical role of Diptera in maintaining network robustness for *M. senegalensis* and the significant trade-offs between flower production and nectar sugar concentration. Flower production, soil moisture and conspecific flowering neighborhood negatively influenced the reproductive outcomes, suggesting the role of pollinator-mediated competition. Future experimental studies could test if such competition could lead to further diversification of the existing interaction niches. Overall, these findings provide valuable insights into the mechanisms driving individual-level plant-pollinator interactions and the importance of preserving keystone individuals to support ecosystem resilience in resource-limited environments.

## Declarations

### Ethics approval and consent to participate

This work did not require ethical approval.

## Consent for publication

Not applicable.

## Availability of data and materials

Datasets will be available from the corresponding author on request.

## Author contributions

D.M.: Methodology, Data curation, Formal analysis, Investigation, Visualisation, Writing-Original draft, Writing-review & editing. K.J.: Investigation, Writing-review & editing. S.K.: Methodology, Conceptualization, Formal analysis, Funding acquisition, Investigation, Supervision, Writing-review & editing.

