## Supplementary material for "Individuals matter: habitat factors and plant traits shape individual-level pollinator interactions in a semi-arid landscape"

### 1. Supplementary Methods

##### 1.1 Spatial distribution of individuals

Based on flowering availability, 31 *Maytenus* individuals were selected for the study. Around each focal individual, a 10 × 10 m plot was established. Within each plot, we recorded the number, identity, flowering phenological status, and distance of all neighboring individuals from the focal plant.

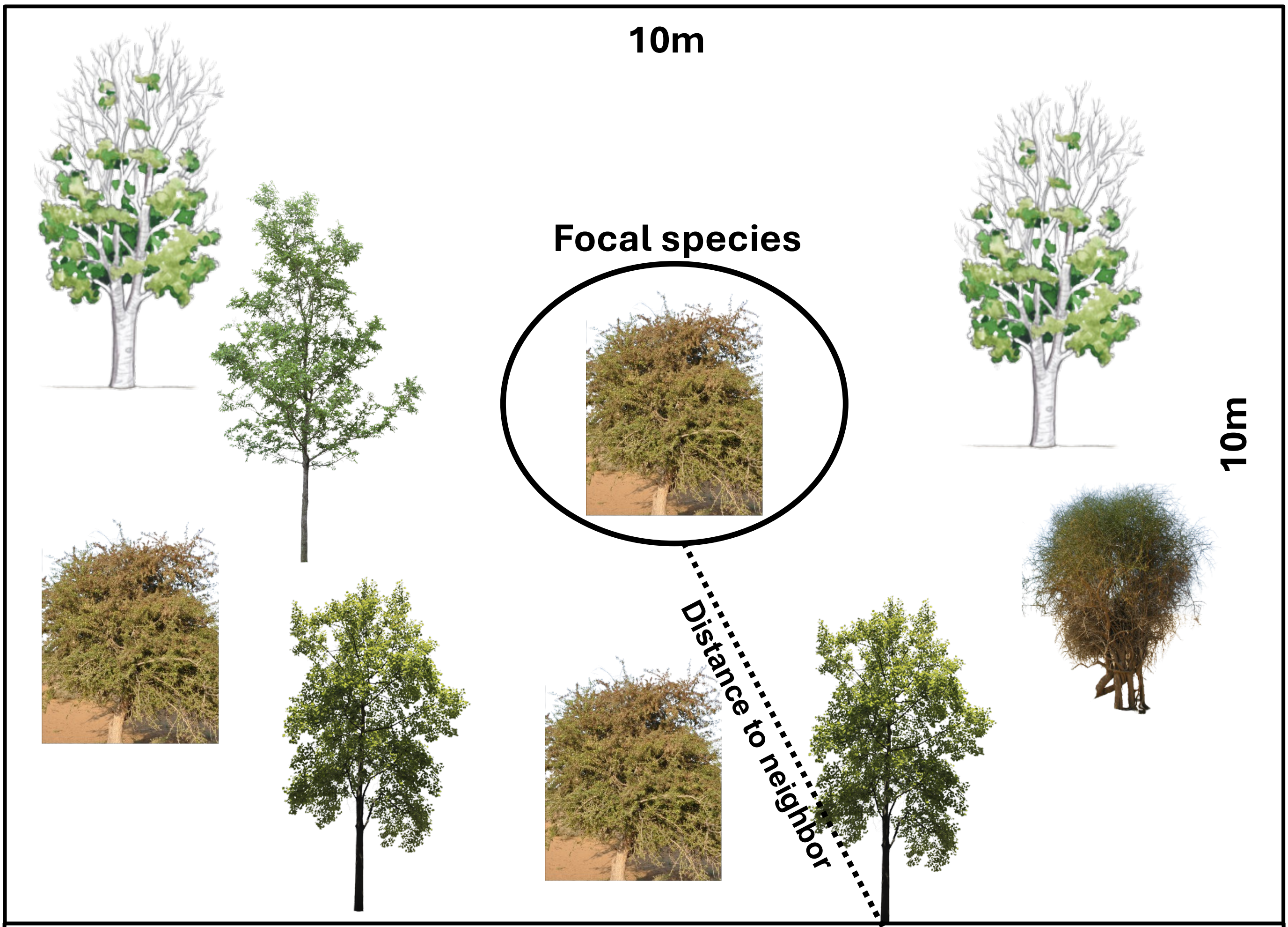


10×10 m plot design

##### 1.2 Spatial autocorrelation test

We performed a spatial autocorrelation test to examine the spatial distribution patterns of individual plants. The test revealed that *Maytenus* individuals exhibited significant positive autocorrelation (Moran's I = 0.502, $p < 0.001$), indicating clustering in soil moisture across the study site. This suggests that individuals occurring in close proximity experienced similar soil moisture levels compared to those located farther apart.

**1.3 Summary of predictor and response variables used in the study, including their sample sizes and the analyses in which each variables are used.**

| Variable | Units | Replicate level | Sample size | Comparison / analysis | Notes / Fig–Table ref. | Remarks |
|---|---|---|---|---|---|---|
| Flower diameter | mm | flowers per plant | 5 flowers/plant; 27 plants | Used in Correlations, ERGM, GLM and GLMMs | **Figures:** 3, 6, 7, S1, S6, S8 **Tables:** 1, 2, S1,S2, S6 | |
| Flower production | count | plant-level (quadrat counts) | all focal plants (sum over visits); 31 plants | Used in Correlations, ERGM, GLM and GLMMs | **Figures:** 3, 6, 7, S1, S6, S8 **Tables:** 1, 2, S1,S2, S6, S7 | |
| Nectar sugar concentration | % Brix | flowers per plant | 8–12 flowers/plant; 28 plants | Used in Correlations, ERGM, GLM and GLMMs | **Figures:** 3, 6, 7, S1, S6, S8 **Tables:** 1, 2, S1,S2, S6 | Nectar concentration measurements for all focal individuals were not possible, due to the low volume in some plants |
| Soil moisture | percentage | plant-level | 4 readings per month and then calculated the average of averages of each month; 31 plants | Used in Correlations, ERGM, GLM and GLMMs | **Figures:** 6, 7, S1, S6, S8 **Tables:** 1, 2, S1, S6, S7 | |
| Distance to habitat edge | m | plant-level | 31 plants | Used in ERGM, GLM and GLMMs | **Figures:** 6, 7, S1, S6, S8 **Tables:** 1, 2, S1, S6, S7 | |
| Proportion of conspecific neighbors | 0-1 | plant-level | No. of conspecifics in a 10×10 m plot / total no. of individuals in the plot; 31 plants | Used in ERGM, GLM and GLMMs | **Figures:** 6, 7, S1, S6, S8 **Tables:** 1, 2, S1, S6, S7 | |
| Pollinator abundance | count / 15-min | 15-min slots per plant | 30–120 min per plant across 34 days; counts per slot | Used in GLM | **Figure** S8 **Table** S8 | |
| Pollinator visits | count / 15-min | 15-min slots per plant | 30–120 min per plant across 34 days; counts per slot | Used in Bipartite network, ERGM and GLM | **Figures:** 4, S8 **Tables:** 1, S8 | |

| Variable | Units | Replicate level | Sample size | Comparison / analysis | Notes / Fig–Table ref. | Remarks |
|---|---|---|---|---|---|---|
| Pollinator identity (morphotypes) | categorical | visit record | 78 morphotypes (48 genus, 7 family, 23 order) | Network (full; No Diptera; No Hymenoptera) | **Figures**: 4, S4 | |
| Conspecific pollen loads | grains per stigma | flowers per plant | 25 flowers/plant; 18 plants | GLM and GLMM vs floral traits/extrinsic factors/Pollinators | **Figures**: 6, S8 **Tables**: 2, S7, S8 | Due to the short flowering period, conspecific pollen loads for a subset of plants was collected |
| Fruit set | % | buds per plant | 10–15 buds/plant; 31 plants | GLM vs traits/extrinsic factors/Pollinators | **Figures**: 7, S8 **Tables**: 2, S7, S8 | |
| H2 (specialization) | 0–1 | network level | 31 plants and 43 pollinator groups | Tested randomness of the network with null networks; Full vs No Diptera vs No Hymenoptera network comparisons; module tests | Degree of specialization in network | |
| Weighted NODF (nestedness) | 0–100 | network level | 31 plants and 43 pollinator groups | Tested randomness of the network with null networks; Full vs No Diptera vs No Hymenoptera network comparisons; module tests | Understands the nestedness in network | |
| Linkage density | >0 = highly connected network | network level | 31 plants and 43 pollinator groups | Tested randomness of the network with null networks; Full vs No Diptera vs No Hymenoptera network comparisons; module tests | Average no. of interactions per individual plant | |
| Shannon diversity | – | network level | 31 plants and 43 pollinator groups | Tested randomness of the network with null networks; Full vs No Diptera vs No Hymenoptera network comparisons; module tests | Interaction diversity by considering both evenness and richness | |

| Variable | Units | Replicate level | Sample size | Comparison / analysis | Notes / Fig–Table ref. | Remarks |
|---|---|---|---|---|---|---|
| Weighted connectance | 0-1 | network level | 31 plants and 43 pollinator groups | Tested randomness of the network with null networks; Full vs No Diptera vs No Hymenoptera network comparisons; module tests | Proportion of realized interactions | |
| PDI | 0-1 | plant-level | 31 plants | Degree of specialization of an individual | **Figure** 5 **Table** S5 | |
| Normalised degree | 0-1 | plant-level | 31 plants | Degree of individual plants as proportion of the possible degree | **Figure** 5 **Table** S5 | |
| Weighted closeness | – | plant-level | 31 plants | Efficiency in which one plant connects to other plant through shared pollinators | **Figure** 5 **Table** S5 | |
| Niche overlap | 0-1 | plant-level | 31 plants | Measures extent to which a plant shares pollinator resources with other plants in the network | **Figure** 5 **Table** S5 | |

#### 2. Supplementary Results

**Table S1** Relationships between soil moisture and three floral traits quantified by Spearman's rank correlation (cor = correlation coefficient). Significant correlations are in boldface ($p < 0.05$). Soil moisture % and flower production data are from 31 plants, whereas nectar concentration and flower diameter (mm) data are from 27 and 28 plants, respectively.

| Trait 1 | Trait 2 | cor | *p*-value | Method |
|---|---|---|---|---|
| Soil moisture | Flower diameter | 0.27 | **0.001** | Spearman |
| Soil moisture | Nectar concentration | -0.02 | 0.69 | Spearman |
| Soil moisture | Flower production | 0.2 | 0.27 | Spearman |

**Table S2** Relationships among the three floral traits quantified by Pearson correlation (cor = correlation coefficient). Significant correlations are in boldface ($p < 0.05$). Flower production data are from 31 plants, whereas nectar concentration and flower diameter (mm) data are from 27 and 28 plants, respectively.

| Trait 1 | Trait 2 | cor | *p*-value |
|---|---|---|---|

| | | | |
|---|---|---|---|
| Flower diameter | log (Flower production) | 0.11 | 0.55 |
| Nectar sugar concentration | log (Flower production) | -0.49 | **0.007** |
| Nectar sugar concentration | Flower diameter | -0.16 | 0.42 |

**Table S3** Table showing the *z*-scores of the network metrics between the observed and randomly generated networks.

| Network metrics | *z*-scores |
|---|---|
| Weighted NODF[1] | -10.87 |
| Weighted connectance[2] | -28.01 |
| Shannon diversity | -9.7 |
| Linkage density[2] | -28.23 |
| log (H2) | 29.53 |

**Table S4** Table showing the percentage of difference in the network metrics between the full network and the network without Diptera as well as the network without Hymenoptera.

| Network-level metrics | % difference between full network and network-without Diptera | % difference between full network and network-without Hymenoptera |
|---|---|---|
| H2 | 36.5 | 2.33 |
| Shannon diversity | 15.1 | 3.12 |
| Linkage density | 27.9 | 9.68 |
| Weighted NODF | 22.8 | 3.35 |
| Weighted connectance | 10.4 | 26.8 |

**Table S5** Table showing the mean, standard deviation and range of the individual plant level metrics ($n$ = 31 plants).

| Node-metrics | Mean | SD | Range |
|---|---|---|---|
| PDI[3,4] | 0.95 | 0.01 | 0.90 - 0.99 |
| Weighted closeness[5] | 0.02 | 0.02 | 0.005 - 0.09 |
| Normalised degree[5] | 0.27 | 0.07 | 0.09 - 0.41 |
| Niche overlap | 0.52 | 0.15 | 0.16 – 0.67 |
| Species strength[6] | 1.38 | 0.88 | 0.12 - 3.53 |

**Table S6** Values of the Spearman correlation coefficients (cor) for the three floral traits with species strength and niche overlap. Flower production data are from 31 plants, whereas nectar concentration and flower diameter data are from 27 and 28 plants, respectively.

| **Trait 1** | **Trait 2** | **cor** | ***p-value*** |
|---|---|---|---|
| Nectar concentration | Species strength | 0.18 | 0.34 |
| Flower diameter | Species strength | -0.09 | 0.63 |
| Flower production | Species strength | -0.08 | 0.65 |
| Nectar concentration | Niche overlap | -0.01 | 0.92 |
| Flower diameter | Niche overlap | 0.19 | 0.33 |
| Flower production | Niche overlap | -0.02 | 0.89 |

**Table S7** Results of the generalised linear models (GLM) evaluating the effects of the flower production and habitat factors of plant individuals on conspecific pollen loads (median, $n = 18$ plants). Response variable was log-transformed and gaussian family was used in the models. Significant values $p < 0.05$ appear in boldface.

| Response | Predictor(s) | Estimate | Std. Error | *z*-value | *p*-value |
|---|---|---|---|---|---|
| Conspecific pollen loads (Model 5) | Soil moisture | 0.231 | 0.101 | 2.281 | **0.022** |
| | Distance to habitat edge | $0.7\times10^{-4}$ | 0.001 | 0.063 | 0.950 |
| | Proportion of conspecific neighbors | -1.599 | 0.561 | -2.847 | **0.004** |
| Conspecific pollen loads (Model 6) | log(Flower production) | -0.694 | 0.352 | -1.967 | **0.049** |

**Table S8** GLM analysis of reproductive success (fruit set %; $n = 31$ plants and conspecific pollen loads; $n = 18$ plants) in response to the abundance of pollinators per minute and number of pollinator visits per minute. Both response variables were log-transformed, and the Gaussian family was used for all models. Significant values $p < 0.05$ appear in boldface.

| **Model No.** | **Response variables** | **Predictor variables** | **Estimate** | **Std. Error** | ***z*-value** | ***p*-value** |
|---|---|---|---|---|---|---|
| Model 7 | Fruit set | Number of pollinators | 0.256 | 0.147 | 1.731 | 0.083 |
| Model 8 | Fruit set | Number of pollinator visits | 0.098 | 0.041 | 2.367 | **0.017** |
| Model 9 | Conspecific pollen loads | Number of pollinators | -0.219 | 0.122 | -1.791 | 0.073 |
| Model 10 | Conspecific pollen loads | Number of pollinator visits | -0.001 | 0.038 | -0.027 | 0.978 |

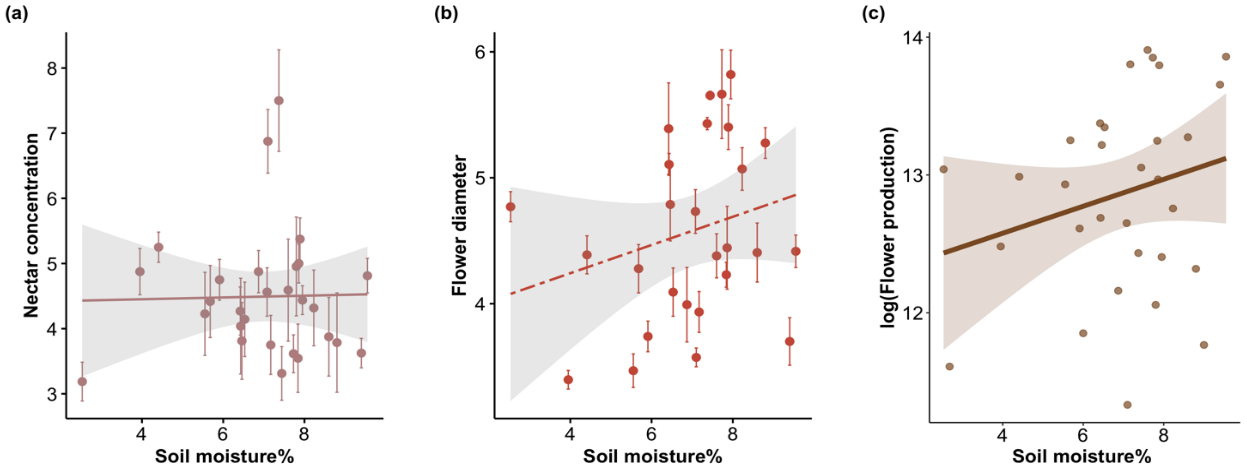


**Fig. S1** Scatter plots illustrating the relationship between soil moisture percentage and the floral traits of individual plants. Floral traits include (a) nectar sugar concentration (% Brix), (b) flower diameter (mm), and (c) flower production ($n$ = 31 plants). Each point represents the mean and ± SE of an individual plant in case of flower diameter and nectar concentration. Dashed fit line indicates significant effect. Nectar concentration and flower diameter data are from 27 and 28 plants, respectively.

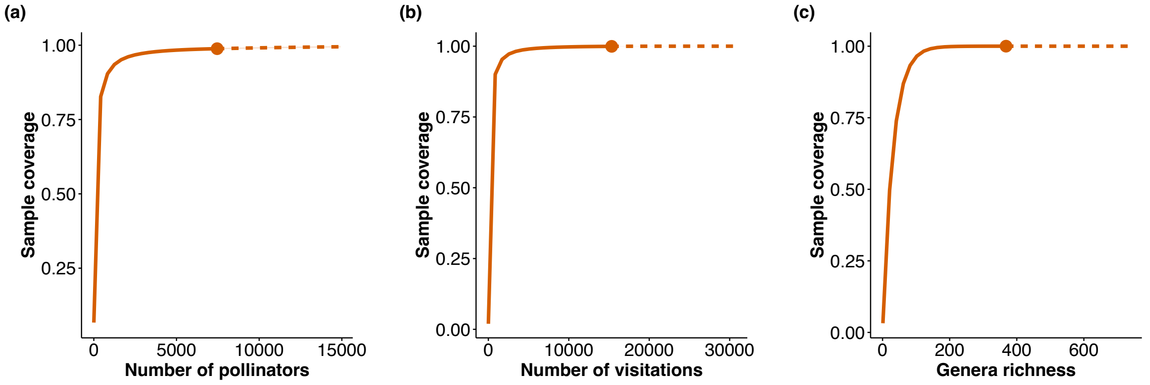


**Fig. S2** Sampling completeness analysis showing (a) the cumulative number of pollinator species, (b) the cumulative number of pollinator visitations, and (c) genera richness across sampling efforts.

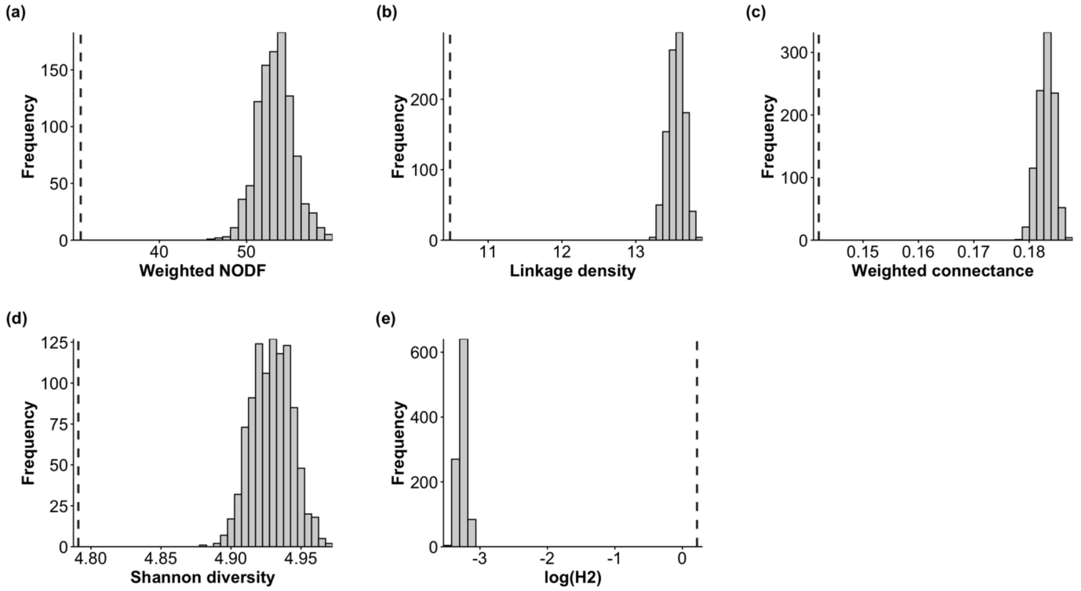


**Fig. S3** Bar graphs displaying the distribution of network metric values from randomly generated networks. The bars represent the frequency of random networks with similar metric values, while the dashed line indicates the corresponding metric value from the observed network.

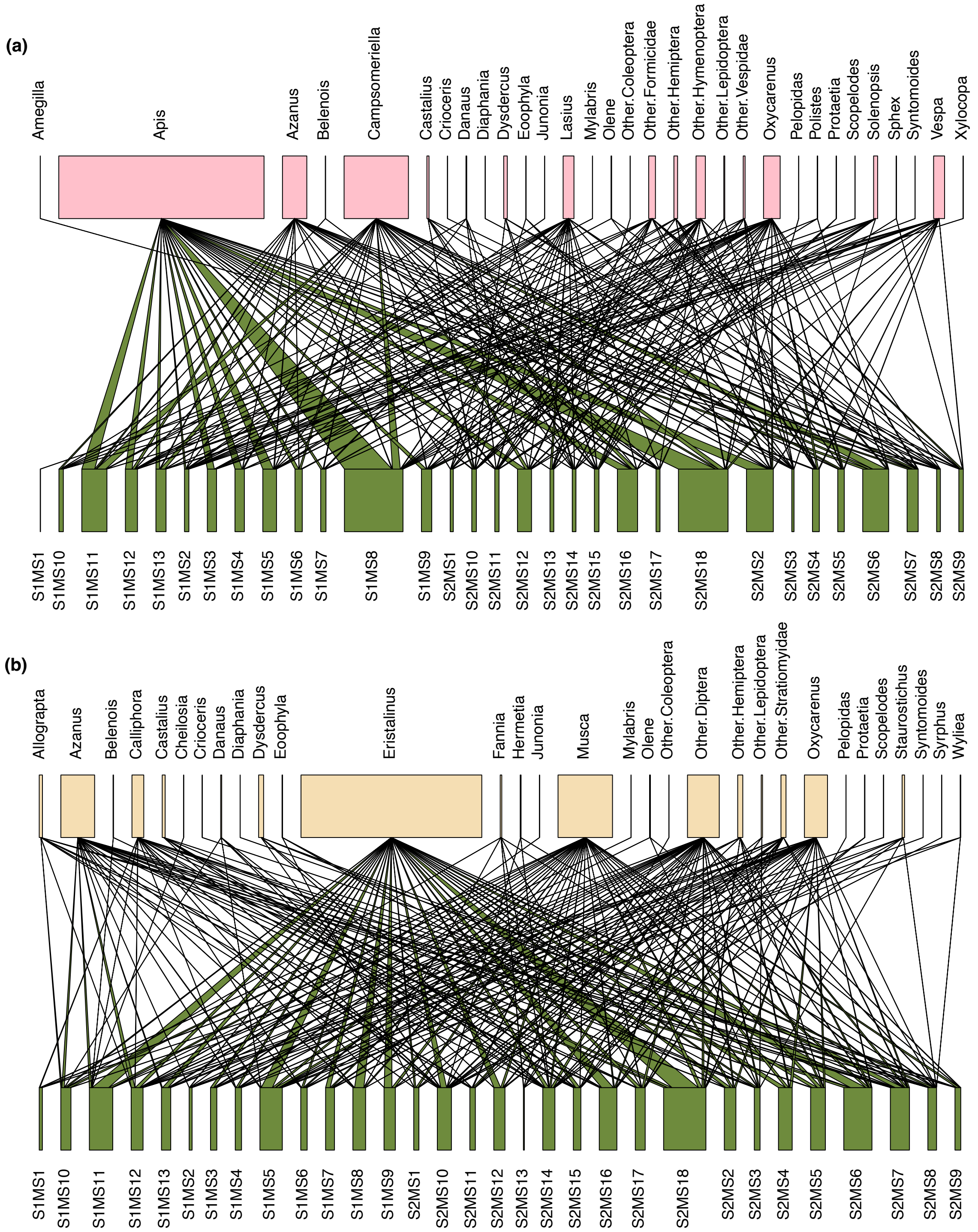


**Fig. S4** Modified weighted bipartite networks by removing the dominant pollinator groups. (a) Network without Diptera: showing the interactions between pollinator species (pink) and *M. senegalensis* individuals (green). (b) Network without Hymenoptera: showing the interactions between pollinator species (light yellow) and *M. senegalensis* individuals (green). The links between nodes represent flower visitation interactions, with the width of the links corresponding to the number of visitations recorded. The pollinators mentioned as ‘Other’ are the morphotypes that could not be identified until the genus level (eg: Other Coleoptera).

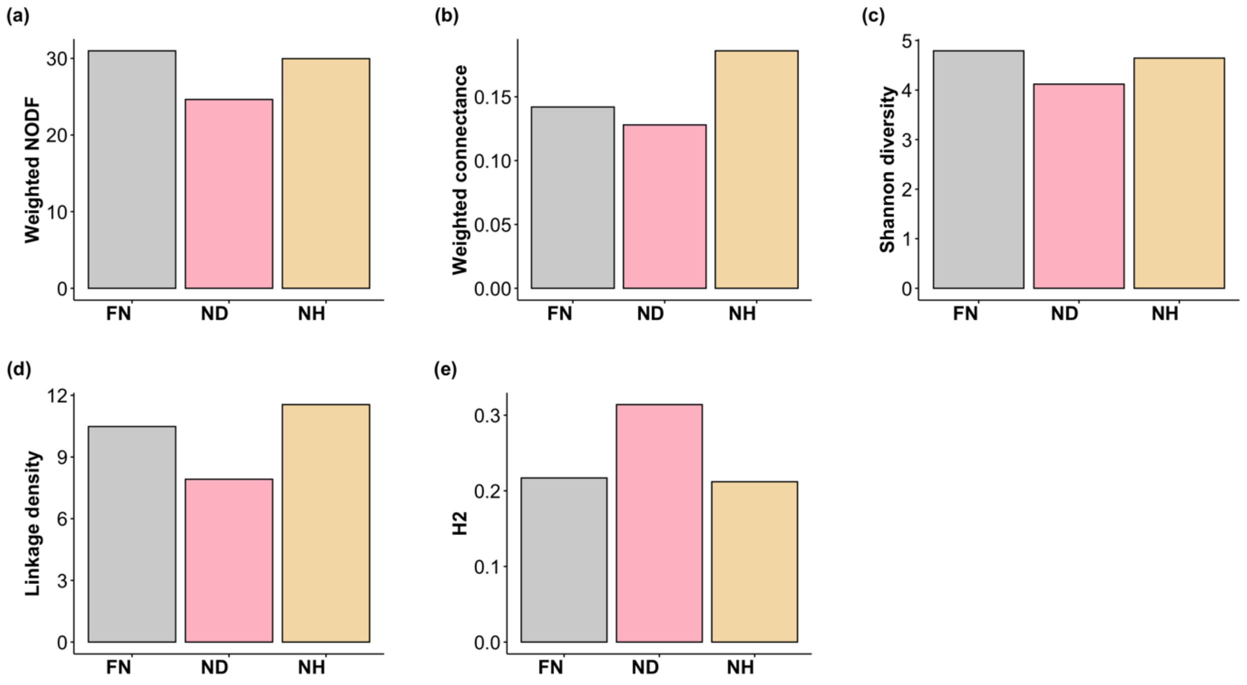


**Fig. S5** Bar graphs illustrating the variation in network metrics across different bipartite networks. The bars represent the following networks: grey - observed network, pink - network without Diptera, and light yellow - network without Hymenoptera.

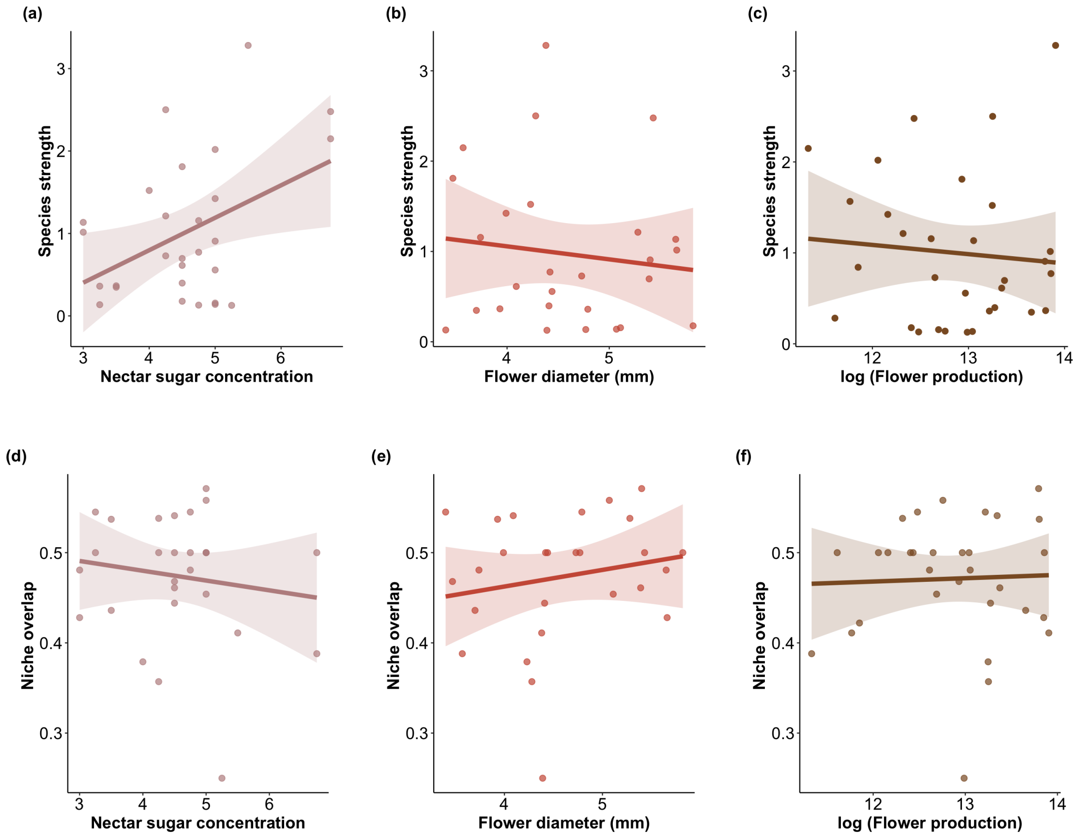


**Fig S6** Scatter plots depicting the variability in species strength of individual plants ($n$ = 31 plants), explained by (a) nectar sugar concentration (% Brix), (b) flower diameter (mm), and (c) flower production; variability in niche overlap (binary; Schoener's index) of individual plants, explained by (d) nectar sugar concentration (% Brix), (e) flower diameter, and (f) flower production. Each point represents an individual plant.

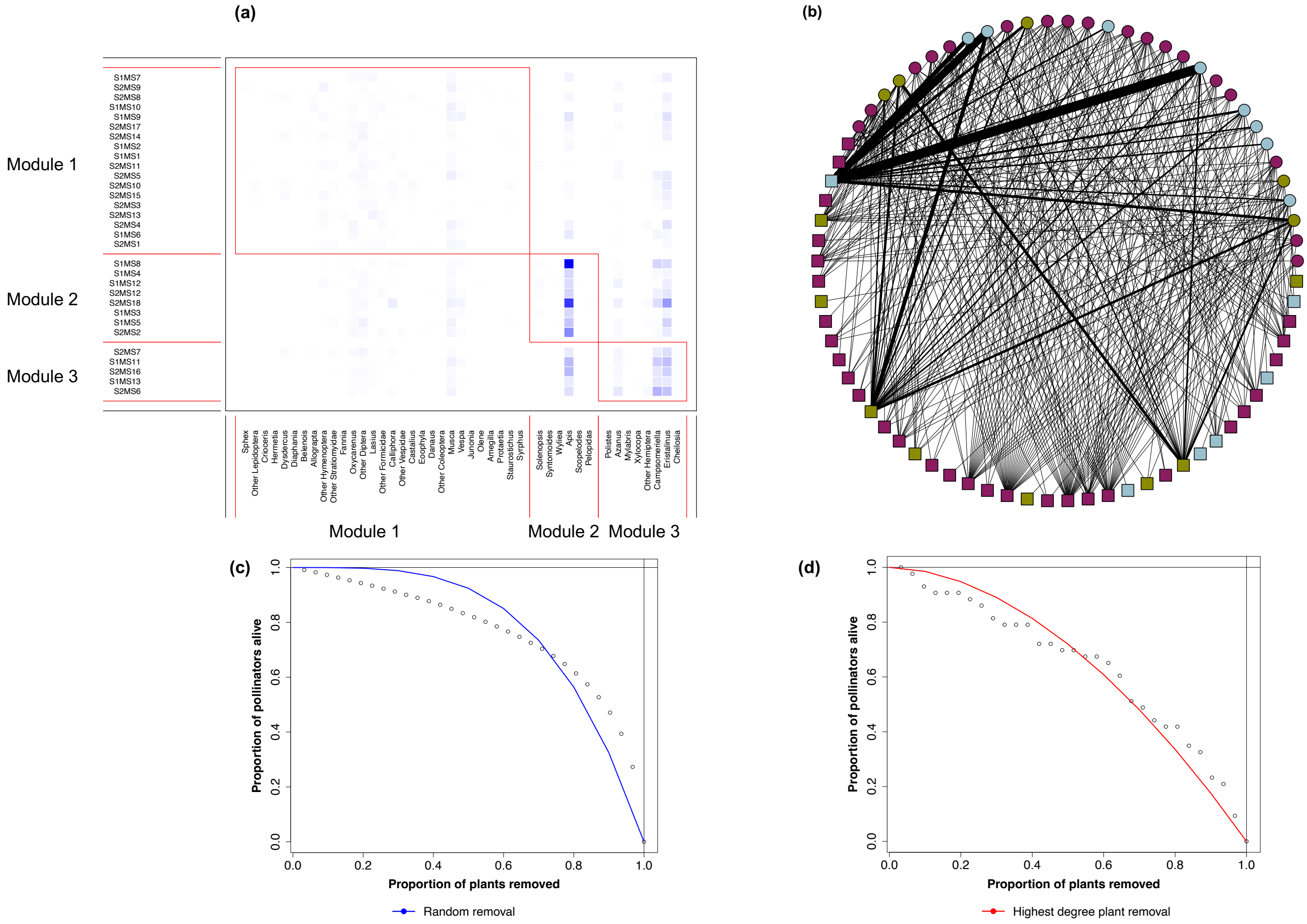


**Fig. S7** (a) Network modularity structure showing modules with pollinators on the x-axis and plant individuals on the y-axis. (b) Visualization of plant modules, where squares represent pollinators, circles represent plant individuals, and different colors indicate distinct modules within the network. (c) The impact of random plant removal on the proportion of remaining pollinators. (d) The effect of sequential removal of plants with the highest degree on the proportion of remaining pollinators (each circle represents an individual plant).

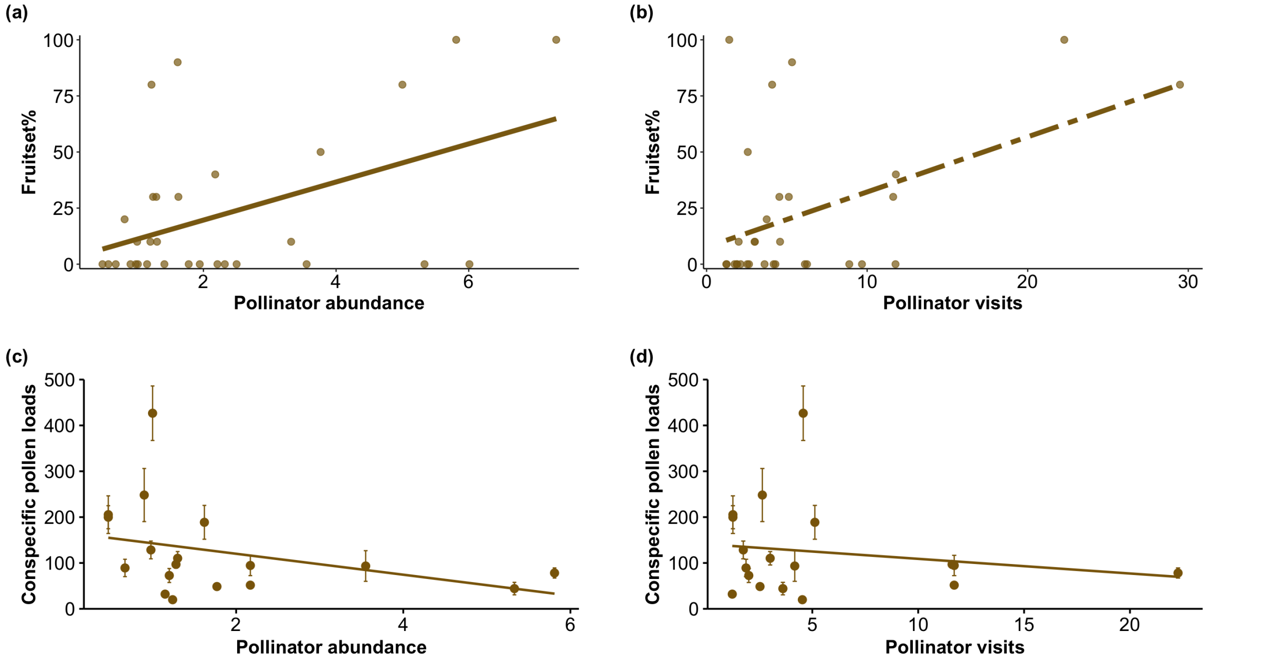

**Fig. S8** Scatter plots depicting the variability in the reproductive success (fruit set %: $n = 31$ plants; conspecific pollen loads: $n = 18$ plants) of individual plants, explained by (a) abundance of pollinators per minute, (b) number of pollinator visits per minute, (c) abundance of pollinators per minute and (d) number of pollinator visits per minute. Each point represents an individual plant. Mean and ± SE has been shown in the plots with conspecific pollen loads as response.

#### 3. Details of statistical analyses

##### ERGM model development to assess the effects of various factors on plant-pollinator interactions

To investigate the effects of floral traits, microhabitat factors, and spatial neighborhood on plant-pollinator interactions, we used binary Exponential Random Graph Models (ERGMs)[7].

**Model Specification**

Each model was constructed based on specific hypotheses regarding the influence of predictor variables. Interaction terms were not included to maintain model simplicity and improve model fit. Additionally, log transformations were applied to certain variables to meet model assumptions and enhance fit.

**Effect of floral traits on plant-pollinator interactions**

Given our hypothesis that intraspecific variation in floral traits influences plant-pollinator interactions, we modelled incorporating floral traits (nectar concentration, flower diameter) as node factors.

Model 1: Effect of floral traits on plant-pollinator interactions
*Edges ~ nodemain(Nectar concentration) + nodemain(Flower diameter)*

**Combined effect of microhabitat and floral traits on plant-pollinator interactions**

To examine the role of microhabitat conditions, we included soil moisture, and to examine the effect of floral traits, we included flower production into this model, hypothesizing that these factors might influence plant-pollinator interactions.

Model 2: Effect of soil moisture and flower production on plant-pollinator interactions
*Edges ~ nodemain(flower production) + nodemain(soil moisture)*

**Combined effect of spatial neighborhood and habitat factors on plant-pollinator interactions**

To understand how spatial neighborhood dynamics affect plant-pollinator interactions, we included proportion of conspecific neighbors, and the distance to habitat edge in the model.

Model 3: Effect of proportion of conspecific neighbors and distance to habitat edge on plant-pollinator interactions

*Edges ~ nodemain(Proportion of conspecific neighbors) + nodemain(Distance to habitat edge)*

##### GLM model development to assess the effects of various factors on reproductive success

To investigate the effects of floral traits, habitat factors, and spatial neighborhood on reproductive success, we used multiple Generalized Linear Models (GLMs). The response variables were fruit set and conspecific pollen loads.

**Model Specification**

Each model was constructed based on specific hypotheses regarding the influence of predictor variables. Interaction terms were checked and not included for final analyses as there were no significant effect of interactions. Additionally, log transformations were applied to certain variables to meet model assumptions and enhance fit.

##### Effect of Floral Traits on Reproductive Success

Given our hypothesis that intraspecific variation in floral traits influences reproductive success, we developed models incorporating key floral traits (nectar concentration, flower production, and flower diameter) as predictor variables.

Model 1: Effect of floral traits on fruit set
*glmmTMB(log(Fruitset +2) ~ Nectar concentration + log(Flower production) + Flower diameter*

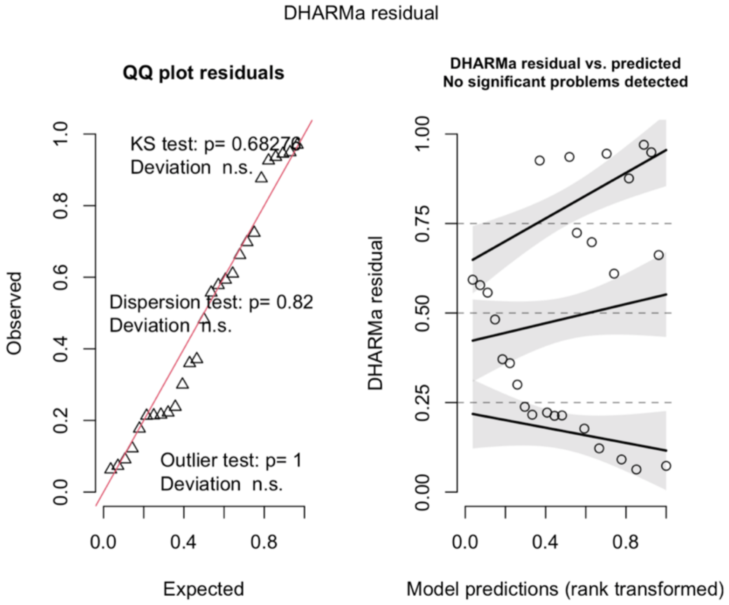


##### Effect of Habitat Factors and spatial neighborhood on Reproductive Success

To examine the role of microhabitat conditions, we included soil moisture, distance to habitat edge and proportion of conspecific neighbors as predictor variables, hypothesizing that these factors might influence both reproductive outcomes.

Model 2: Effect of habitat factors on fruit set
*glmmTMB(log(Fruitset +2) ~ soil moisture + distance to habitat edge + proportion of conspecific neighbors*

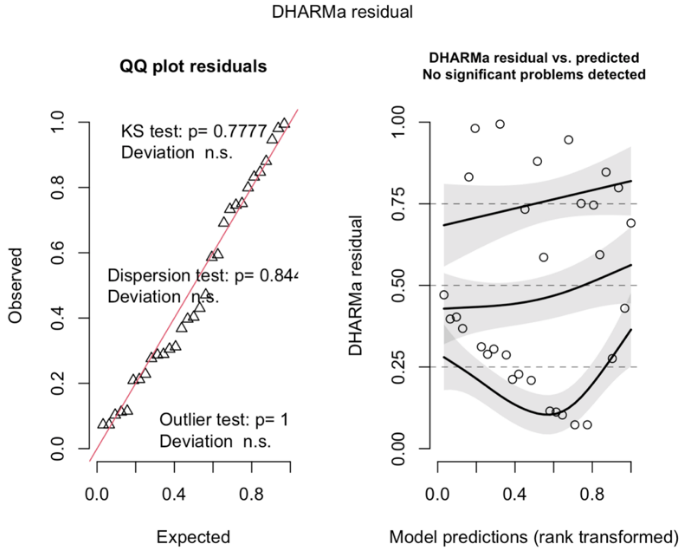

**GLMM model development to assess the effects of various factors on conspecific pollen loads**

To investigate the effects of floral traits, microhabitat conditions, and individual-level variability on conspecific pollen loads, we used multiple Generalized Linear Mixed Models (GLMMs)[8]. In all models, plant individual identity (Plant ID) was included as a random effect to account for the effect of plant individual (within plant variation) on conspecific pollen loads. Predictor variables were selected based on our hypotheses, and log transformations were applied where necessary to improve model fit. Interaction terms were not included to maintain model simplicity and ensure better convergence.

**Effect of Microhabitat Factors on Conspecific Pollen Loads**

To evaluate the role of habitat factors, we included soil moisture, proportion of conspecific neighbors, and distance to habitat edge as fixed effects.

Model 3: Effect of habitat factors and spatial neighborhood

*glmmTMB (log(Conspecific pollen loads) ~ soil moisture + distance to habitat edge+ proportion of conspecific neighbors + (1/Plant ID)*

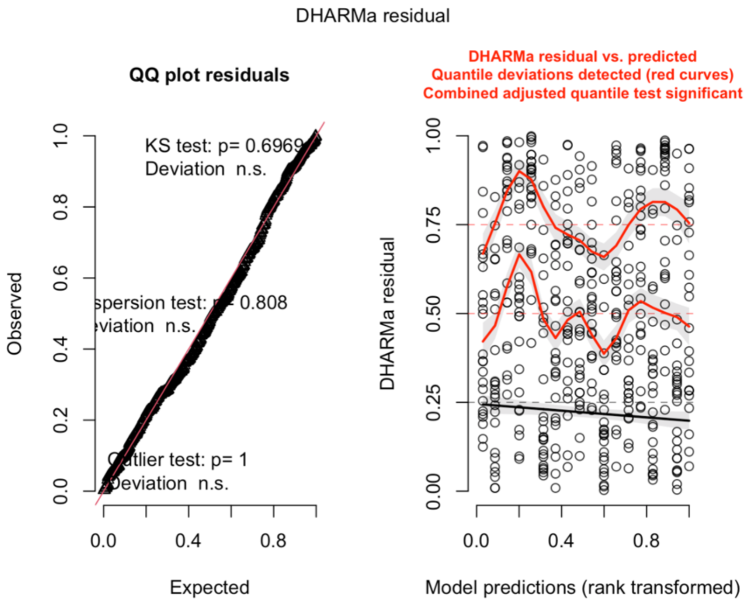


**Effect of Floral Traits on Conspecific Pollen Loads**

This model assessed the influence of key floral traits on conspecific pollen loads. We included flower production, nectar concentration and flower diameter as fixed effects. This model aimed to evaluate how this combination of floral traits influences conspecific pollen deposition.

Model 4: Effect of floral traits on pollen loads

*glmmTMB (logConspecific pollen loads ~ scale(Nectar concentration) + scale(Flower diameter) + log(Flower production) + (1/Plant ID)*

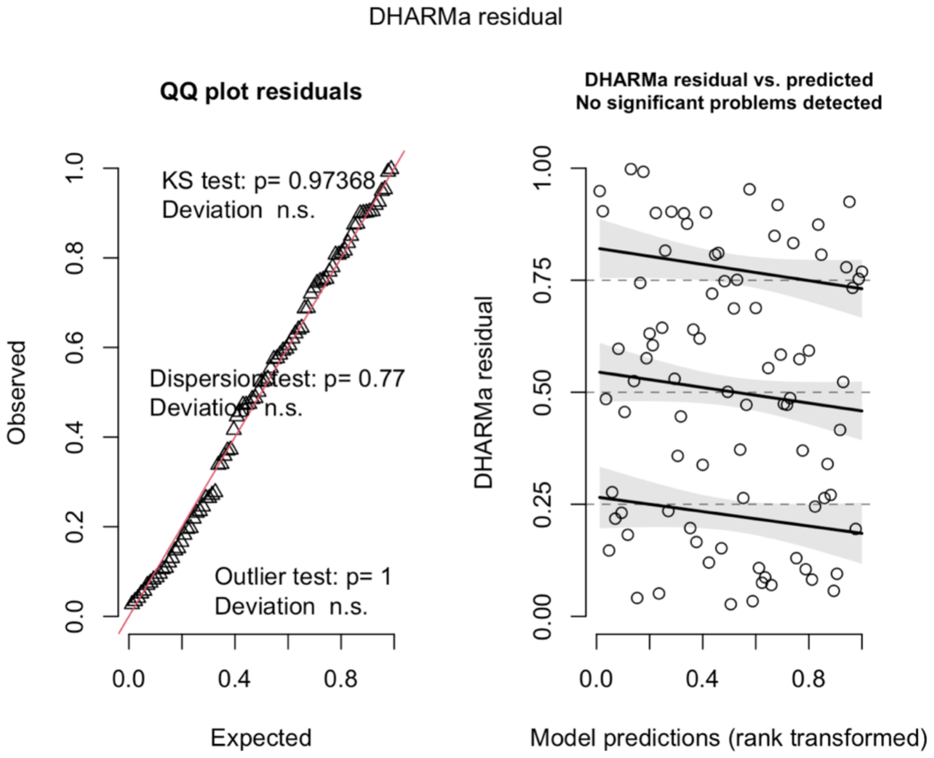


Fixed effects included in these models such as soil moisture, flower diameter, and proportion of conspecific neighbors significantly influenced conspecific pollen loads. Apart from that, the random effect of individual plants also accounted for a substantial proportion of the variability in conspecific pollen loads, indicating that plant-specific factors not captured by the fixed effects play a role in determining conspecific pollen deposition (Fig. 6, Table S7).

**Goodness of fit plots for models 5-10 described in Table S7 and S8**

**Model 5**

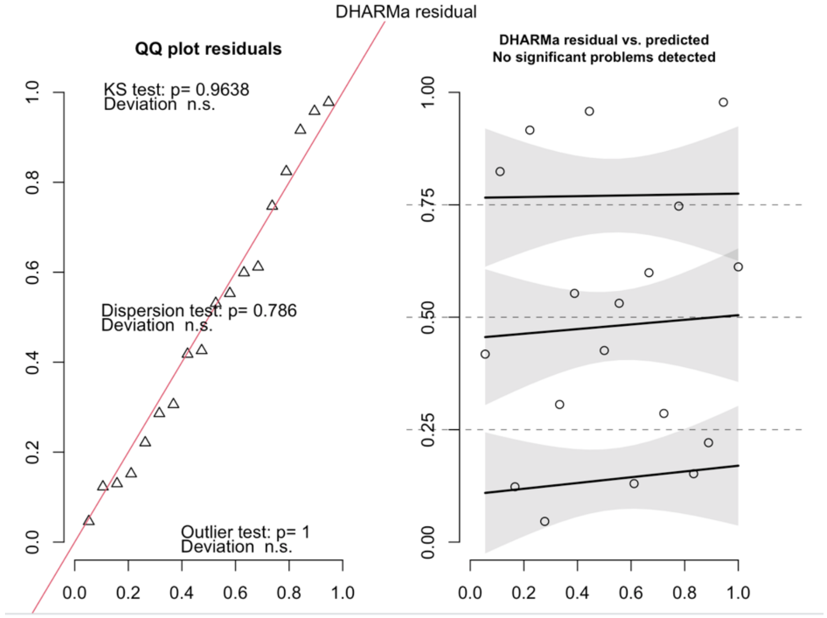

**Model 6**

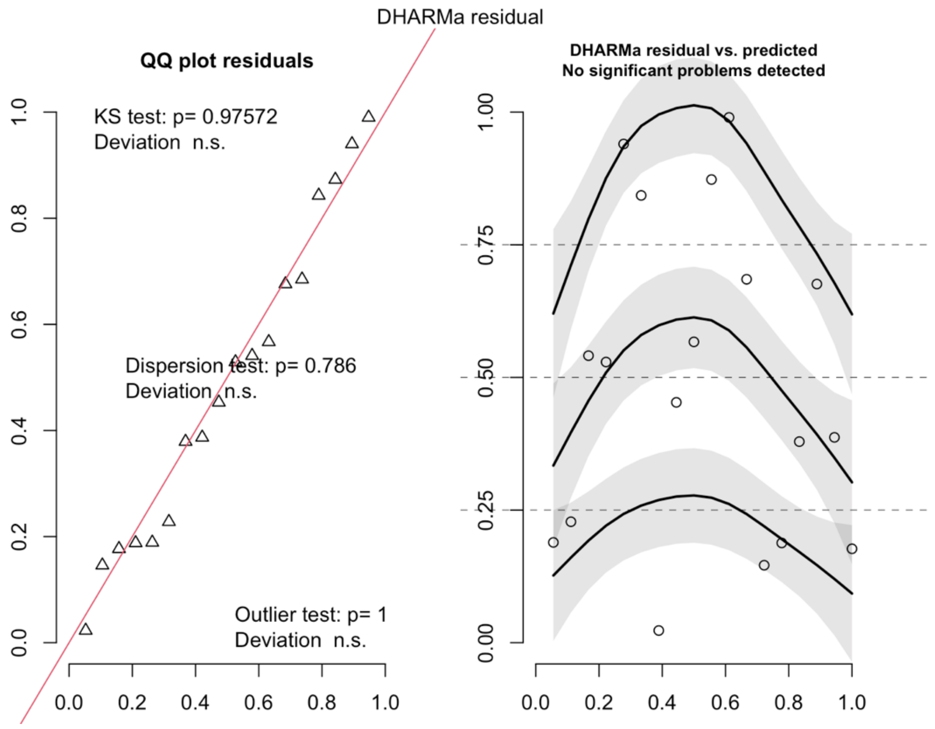


**Model 7**

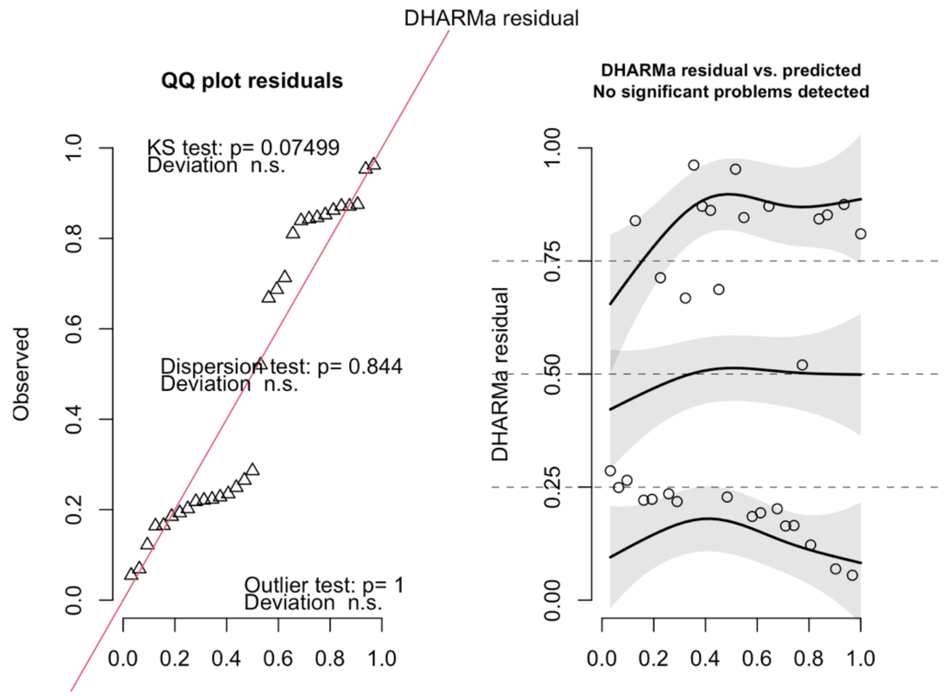

**Model 8**

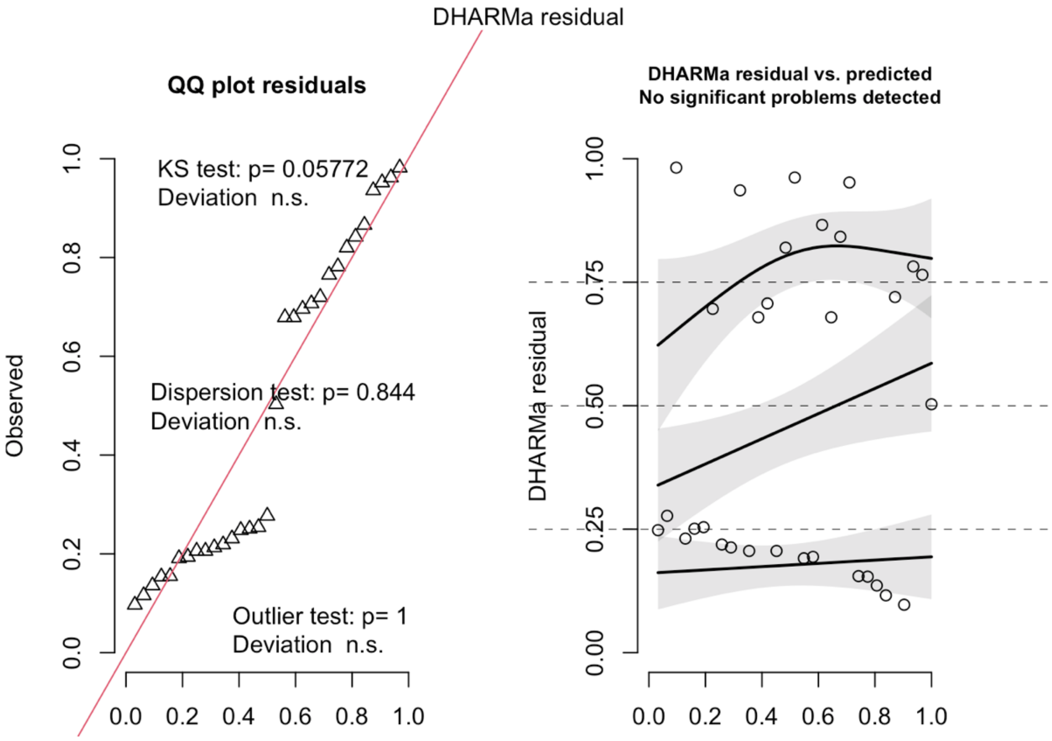


**Model 9**

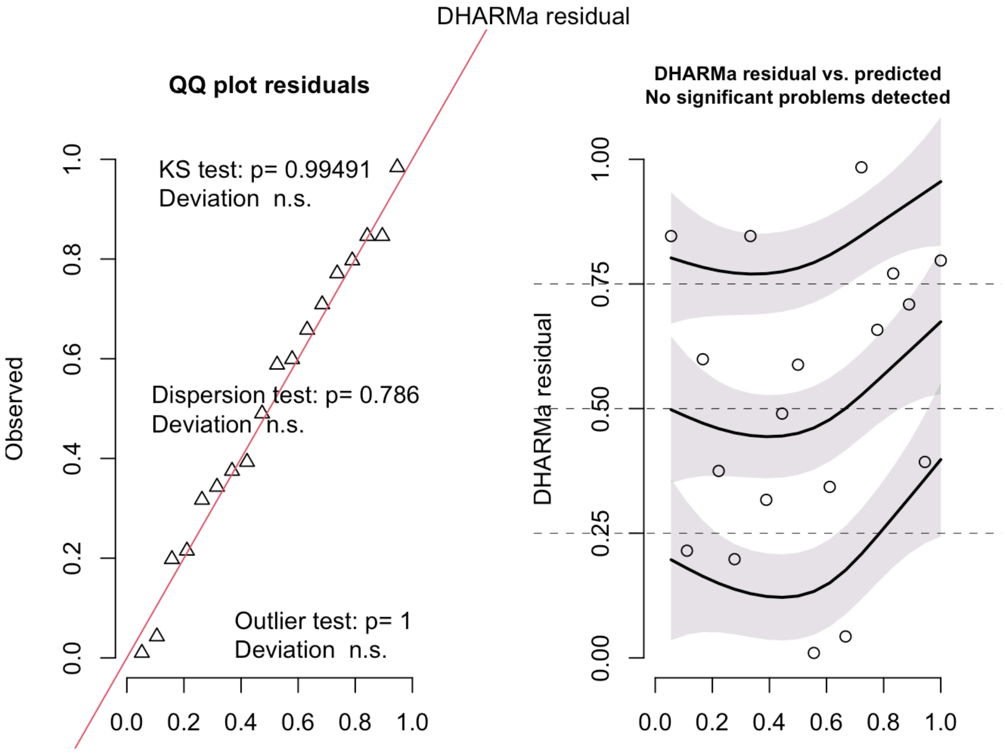

**Model 10**

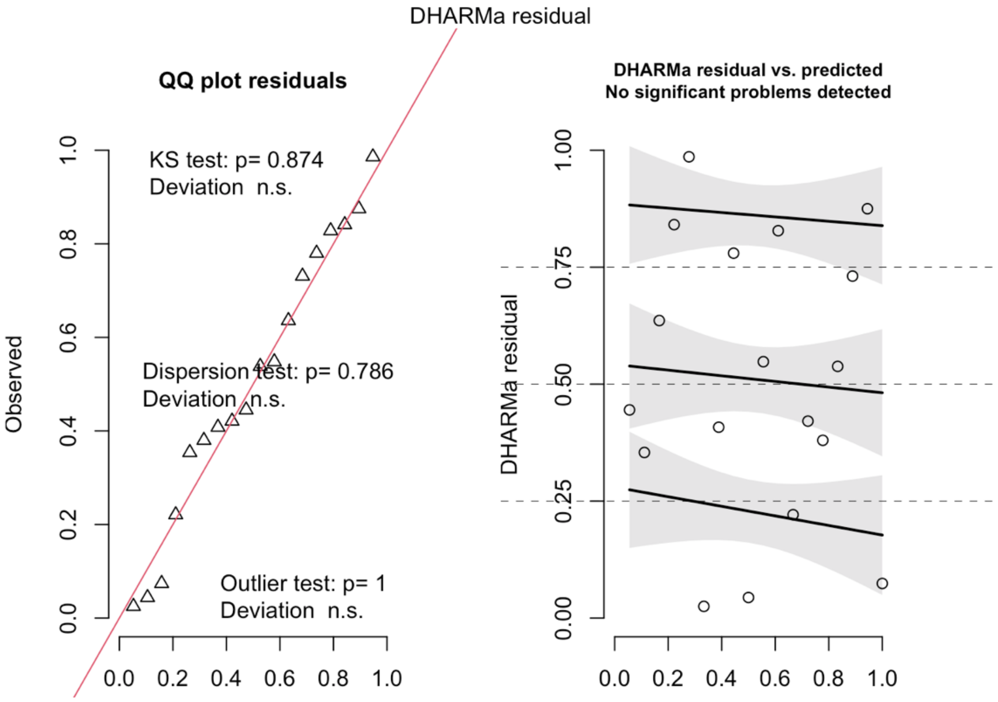
